# Distinct gene programs underpinning ‘disease tolerance’ and ‘resistance’ in influenza virus infection

**DOI:** 10.1101/2022.03.22.485254

**Authors:** Ofir Cohn, Gal Yankovitz, Naama Peshes-Yaloz, Yael Steuerman, Amit Frishberg, Rachel Brandes, Michal Mandelboim, Jennifer R. Hamilton, Tzachi Hagai, Ido Amit, Mihai G. Netea, Nir Hacohen, Fuad A. Iraqi, Eran Bacharach, Irit Gat-Viks

**Affiliations:** The Shmunis School of Biomedicine and Cancer Research, George S. Wise Faculty of Life Sciences, Tel Aviv University, Tel Aviv, Israel; Central Virology Laboratory, Ministry of Health, Chaim Sheba Medical Center, Tel-Hashomer, Ramat Gan, Israel; Department of Microbiology, Icahn School of Medicine at Mount Sinai, New York; Department of Immunology, Weizmann Institute, Rehovot, Israel; Department of Internal Medicine and Radboud Center for Infectious Diseases, Radboud University Medical Center, Nijmegen, the Netherlands; Department for Genomics & Immunoregulation, Life and Medical Sciences Institute (LIMES), University of Bonn, Bonn, Germany; Broad Institute of MIT and Harvard, Cambridge, MA, USA; Massachusetts General Hospital Cancer Center, Boston, MA, USA; Department of Epidemiology and Preventive Medicine, School of Public Health, Sackler Faculty of Medicine, Tel-Aviv University, Tel-Aviv, Israel; Department of Molecular and Cell Biology, University of California, Berkeley, Berkeley, CA 94720, USA

**Author notes:** Corresponding author, (EB), (IGV). These authors contributed equally to this work (Ofir Cohn, Gal Yankovitz). These authors contributed equally to this work (Eran Bacharach, Irit Gat-Viks).

## Abstract

When challenged with an invading pathogen, the host defense response is engaged to eliminate the pathogen (resistance) and to maintain health in the presence of the pathogen (disease tolerance). However, the identification of distinct molecular programs underpinning disease tolerance and resistance remained obscure. We exploited transcriptional and physiological monitoring across 33 mouse strains, during *in vivo* influenza virus infection, to identify two host-defense gene programs – one is associated with hallmarks of disease tolerance and the other with hallmarks of resistance. Both programs constitute generic responses in multiple mouse and human cell types. Our study describes the organizational principles of these programs and validates Arhgdia as a regulator of disease-tolerance states in epithelial cells. We further reveal that the baseline disease-tolerance state in macrophages is associated with the pathophysiological response to injury and infection. Our framework provides a paradigm for the understanding of disease tolerance and resistance at the molecular level.

## Introduction

The host defense against infectious diseases involves two main strategies: ‘disease tolerance’ and ‘resistance’. The resistance strategy enables the host to detect and eliminate the invading pathogen. The ‘disease tolerance’ strategy enables the host to maintain health in the presence of pathogens^1–7^. We note that the conventional terminology of ‘disease tolerance’ should not be confused with ‘immune tolerance’ used to describe the lack of responsiveness to particular antigens^6,8^.

As a critical component of the host defense, disease tolerance has become of particular interest in the treatment of infectious diseases, such as the influenza virus and SARS-CoV-2 infections^1,9–13^. However, a major barrier for the research on disease tolerance is that it typically requires analyses at the physiological and organismal levels, while missing an experimental paradigm for investigations at the cellular and molecular levels; this contrasts with the resistance strategy that is studied at all levels – the molecular, cellular, physiological, and organismal levels (recently reviewed in Refs. 9,10,14). Consequently, current efforts are mainly focused on developing drugs against the pathogen rather than drugs that modulate the host’s ability to promote health in the presence of the pathogen^9,11^. Therefore, there is an urgent need to develop methodologies for the investigation of disease tolerance at the molecular level.

Determining molecular states underpinning disease tolerance remained a challenge due to the close interrelations between disease tolerance and resistance: both disease-tolerance and resistance mechanisms are jointly activated during infections and both types of mechanisms have multiple shared genes and pathways. For instance, NFkB signaling has a role in antiviral signaling (a resistance mechanism) as well as a role in tissue repair (disease-tolerance mechanism)^15–18^. Thus, cell states that are associated with disease tolerance have not been defined independently of resistance, limiting our ability to study this central process at the molecular level.

Here we exploited interindividual variation in the host’ to identify central gene programs that act during the infection defense response. We developed and validated the computational model using *in vivo* influenza A virus (IAV) infection of 33 genetically diverse mouse strains that differ in their capacity to resist and tolerate disease, and by integrating human data of multiple isolated cell types (including non-immune, innate and adaptive immune cells). The analysis demonstrated one gene program that is associated with the strategy of disease tolerance and another gene program that is associated with the resistance strategy, both at the functional and the phenotypic levels. In accordance, we refer to these programs as ‘disease tolerance’ and ‘resistance’ programs, respectively. Systematic analysis of human data across various healthy and inflammatory conditions indicated that the two programs are shared across multiple cell types – including non-immune, innate and adaptive immune cells. Relying on this finding, we developed a generic quantitative metric for the molecular levels of the disease tolerance and resistance programs. This scoring scheme allowed us to identify distinct markers for the levels of each program, and subsequently to reveal Arhgdia as a regulator of the disease-tolerance program during the host response to IAV infection. Importantly, the baseline level of disease tolerance in blood macrophages (MFs) is correlated with pathological responses to stress (both injury and infection), suggesting the basal/early disease-tolerance status of MFs as a druggable driver of disease states. Overall, our approach provides a paradigm for the systematic study of disease-tolerance and resistance states and how they malfunction in complex diseases.

## RESULTS AND DISCUSSION

### A longitudinal study of IAV infection across different genetic backgrounds

To characterize the host defense, we collected longitudinal data during *in vivo* IAV infection across 33 genetically distinct mouse strains of the collaborative cross (CC) cohort^19^ (**Figure 1A**; infection was performed using the H1N1 influenza virus, particularly the PR8 virus strain). We recorded lung transcriptomes both in steady-state (before infection) and at multiple time points during infection. We also recorded several phenotypes for these mice, such as the viral burden in the lungs, whole-body weight loss, breathing dysfunction, and expression signatures for tissue damage based on the reduction of marker genes, either cilium markers or markers of non-immune (CD45^−^) cells (**Table S1, Methods;** see validation of the tissue damage signatures in **Figure S1A**). The selected time interval encompasses the initial incubation period between exposure to the virus and the onset of systemic symptoms (3-24h post infection (p.i)), and the acute stage that is characterized by an exponential increase in viral burden, pronounced symptoms, and robust immune responses (24-96h p.i., **Figure S1B**).

**Figure 1.**
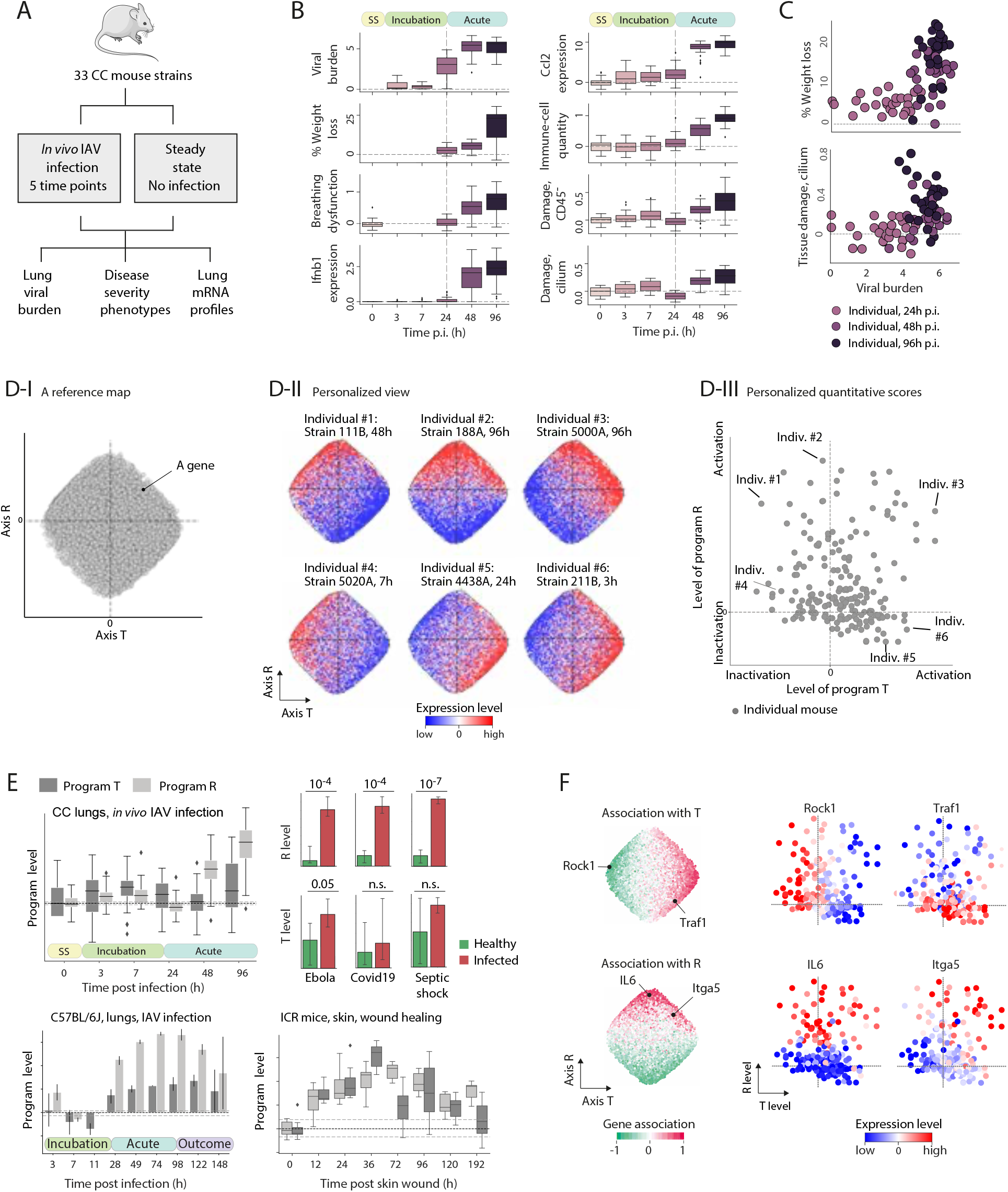
Uncoupling between two key programs of the host response to IAV infection. (**A**) Study design. (**B**) Wide phenotypic diversity of CC mice in response to IAV infection. Each box plot represents diversity across 33 animals of different CC strains, for the indicated phenotype at each time point post infection. SS, steady state. Expression levels are log-scaled. (**C**) Wide phenotypic diversity in the ability to tolerate disease. Presented are the relationship between phenotypes and viral burden across CC individuals (dots), highlighting diversity in disease severity for the same pathogen burden. For comparability, presented are Mx1-deficient CC mice only. (**D**) Definition of gene programs T and R. (I) A gene map. A two-dimensional space in which each gene (a dot) is located in a particular coordinate, such that nearby genes have similar response patterns across all CC individuals. T/R: the horizontal/vertical axes. (II) In each panel, the same map (from D-I) is colored by expression of genes in one selected individual (indicated on top). The blue/red scale indicates low/high expression levels (no data smoothing). For each individual, its gradient along the T and R axes enabled us to calculate scores for the levels of programs T and R, respectively, as shown in plot III. (III) Personalized quantitative scores for the activation levels of programs T and R. Shown are the T/R levels across all CC individuals (dots). Individuals from D-II are indicated. (**E**) Changes of T and R levels in response to several infections and wound healing^53,57, 59–61^. Plots compare the T and R levels before and after stimulations, indicating that increasing T/R levels represent T/R activation. (**F**) Program-associated genes. Left: The map from D-I, where each gene is colored by its association with programs T (top) and R (bottom) (no data smoothing). The association is defined as the Pearson’s correlation between gene expression and program level across all individuals. Right: selected T/R-associated genes. Each indicated gene was colored according to its expression in each individual (a dot), using the landscape of states from D-III; coloring as in D-II.

We found substantial diversity in resistance-related traits, such as viral burden and antiviral responses (**Figure 1B**). Several lines of evidence show that this diversity is not dominated by noise, but rather a biological effect: much of the phenotypic diversity is inherited (**Figure S1C**), there is a significant difference between Mx1-deficient and wild-type CC strains, as expected^20^ (**Figure S1D**), and finally, the antiviral response is largely coordinated with the viral burden, as expected (**Figure S1E**). We also observed substantial differences in the ability to tolerate disease. For instance, as shown in **Figure 1C,** even in the presence of a similar viral burden (viral mRNA expression of around 5, log scaled, at 96h p.i.), some mice lost only 7-10% of their weight (high disease tolerance) while others lost more than 20% of their weight (low disease tolerance). As described below, we followed a stepwise approach to identify key gene programs and to characterize their relations with various hallmarks of disease tolerance and resistance.

### Identification of two central gene programs of the infection defense response

To model the host defense, we used an autoencoder to reduce the multi-dimensionality of the IAV-infection data (i.e., multiple genes, host genetic backgrounds and time points) into a two-dimensional space (**Figure 1D-I, Table S2**). In this arrangement, referred to as a “map”, each gene is embedded at a certain coordinate within the two-dimensional space. The construction of the map relied on a ‘similarity rule’: The closer two genes are to each other in the map, the higher the similarity of their transcriptional responses in all measured strains and at all time points (**Figure S2A**). As we show in subsequent sections, the horizontal and vertical dimensions of the map are linked to disease tolerance and resistance; in accordance, we refer to these axes as “axis T” and “axis R”, respectively.

In many individuals, we observed a gradual change in expression levels over the T and R axes (see color coding of several individuals in **Figures 1D-II, S2B**). Different individuals showed distinct directions and distinct rates of change along their gradients (**Figure 1D-II**), with a nearly linear change in the expression of genes along the gradient (**Figure S2C**). This organization indicates the existence of two regulatory programs, denoted T and R, which underlie the heterogeneity of gene expression along the T and R axes, respectively.

### Personalized quantitative scores for the levels of the two identified programs

To derive quantitative metrics for the activation level of programs T and R in any given individual, we leveraged the gradients of gene expression along the T and R axes. As the gradient of an individual can be decomposed into two gradients that run along the two axes, the personal activation levels of programs T and R can be represented by two quantitative metrics: the T score reflects the gradient along axis T (the ‘activation level of T’) and the R score reflects the gradient along axis R (the ‘activation level of R’) (calculated using a linear deconvolution approach, see **Methods**). Positive and negative activation levels indicate increasing and decreasing gradients along the axes, respectively, and a zero level indicates an absence of a gradient. A two-dimensional representation of individual states by their T and R levels is presented in **Figure 1D-III.** For instance, an individual with a bottom-left to top-right gradient is scored by positive T and R levels (e.g., individual #3 in **Figure 1D-II, III**) and an individual with a top-left to bottom-right gradient is scored by negative T and positive R (e.g., individual #1 in **Figure 1D-II, III**; see additional examples in **Figure S2B**). Computational analysis confirmed that each of the scores is accurate and successfully explains a significant fraction of the variation during infection (**Supplementary Information 1, Figure S3**). The quantitative T and R levels were primarily increasing during inflammation. For example, we observed increased T and R levels during the acute phase of IAV infection, in response to additional pathogens (septic shock, Ebola and SARS-CoV-2 infections), and during wound healing (**Figure 1E;** see data sources in **Methods**). Therefore, the levels of each program scale from inactivation (negative values) to activation (positive values; **Figure 1D-III**).

We used the correlation between the inferred levels of a program and the expression levels of a gene across individuals to determine the association of a gene with a program. The associations provide a continuum of predicted assignments of genes to each program (positive/negative association score indicates a gene that is induced/repressed during activation of a program). As expected, many genes are tightly associated with the programs – for instance, *Il6* is positively-associated with R, but not with T, and Rock1 is negatively associated with T but not with R (**Figures 1F, S4A**). We note that increasing positions of genes in the map (left to right, bottom to top) correspond to increasing associations of genes with the respective programs (**Figures 1F** left and **S4B**). We next use the inferred associations to interrogate the molecular functions of programs T and R.

### The two programs, T and R, are associated with the hallmarks of disease tolerance and resistance, respectively

The host defense strategies have been classified into two broad categories: resistance strategies that sense and react to eradicate the pathogen, and disease-tolerance strategies that enable the host to promote health in the presence of pathogen^1–7^. Comparisons with established hallmarks of disease-tolerance and resistance strategies – including (*i*) molecular functions, (*ii*) response to stressors, and (*iii*) organismal phenotypes – indicated that the T and R programs manifest the main hallmarks of disease-tolerance and resistance, respectively. For each of these aspects, we first describe the methodology and then consider the results.

i. *Molecular functions*. To identify the functional properties of programs T and R, we scored the enrichment of each functional gene set within genes that are associated with each program (a Wilcoxon rank sum test; a positive/negative sign indicates that the activation of the program corresponds to induction/repression of genes annotated with the function; **Methods)**. We collected and analyzed functional categories that have been previously classified as either disease-tolerance (e.g., defensive health functions such as wound healing^2^) or resistance functions (e.g., defensive resistance functions such as antigen presentation). In support of the tolerance/resistance hypothesis, we found that activation of program T was enriched with classified disease-tolerance functions (T: *p* <10^−71^, R: *p* >0.1, *t*-test), and activation of program R was enriched with classified resistance functions (T: *p* >0.1, R: *p* <10^−43^, *t*-test) (**Figure 2A**). For example, the molecular function of wound healing, which has a known disease-tolerance role^2^, is induced during activation of program T but not program R (T-enrichment *q* <10^−12^, R-enrichment *q* >0.05, **Figure 2B, top**). Indeed, *Furin, Pdgfb* and *Tgfb1*, which have known roles in wound healing^21^, are positively correlated with activation of program T (Pearson’s *r =*0.73, 0.56, 0.52, respectively) but not with program R activation (absolute Pearson’s *r* =0.05, −0.12, 0.19, respectively). Another example is ‘cytokine storm syndrome’, which is considered as an uncontrolled resistance response^22^, and in agreement, we found that cytokine storm genes are induced during activation of program R (*q* <10^−6^) but not program T (*q* >0.05, **Figure 2B, bottom**; e.g., *Il1b* and *Tnf*, with Pearson’s *r >*0.72 for R and *r <*0.24 for T). Taken together, the analysis supports the notion that the T and R responses are enriched with a disease-tolerance and resistance functions, respectively. We note that the two programs do not differ based on classical categorizations of immune functions: the two programs involve the interferon and NFkB signaling (**Figure S5A**), the two programs are associated with both pro- and anti-inflammatory genes (**Figure S5B**), and finally, as discussed below, each of the programs acts in a variety of innate and adaptive immune cell types.
ii. *Response to stressors*. The tolerance-resistance paradigm holds that resistance mechanisms primarily responds to biotic stress and that disease-tolerance mechanisms are activated in response to both biotic and abiotic stress^2–4^. To test whether programs T and R fit this paradigm, we calculated enrichments scores for each program using all predefined sets of genes that have a role in various biotic and abiotic stress responses (325 abiotic stress and 30 biotic stress gene sets). As shown in **Figure 2C**, program R is enriched with genes that have a role in various types of biotic stress (*p* <10^−64^) but not abiotic stress (t-test *p*> 0.05), whereas program T is enriched with genes that have a role in both biotic and abiotic stress (*p* <0.003, *p* <0.001). Thus, programs T and R fit the signature of disease-tolerance and resistance mechanisms, respectively (**Figure S5C**).
iii. *Organismal phenotypes*. The previously described tolerance-resistance model postulates that the viral burden is specifically associated with resistance mechanisms but exhibits little to no association with disease-tolerance mechanisms; in contrast, the phenotypic diversity in the ability to tolerate infection (termed a ‘disease tolerance phenotype’) could be affected by both disease-tolerance mechanisms and resistance mechanisms^2,3^. It has been further shown that in some cases, disease tolerance phenotypes are only associated with disease-tolerance mechanisms (i.e., no effect of resistance mechanism^14,23^). To test the relations of programs T and R with organismal phenotypes, we used two cohorts: the CC cohort in this study (infected with H1N1 influenza virus) and an independent CC cohort (infected with H3N2 influenza virus^24^). The phenotype of disease tolerance for each mouse strain was calculated through the standard reaction norms approach^14,25^ (see **Methods** and **Figure S6AB**; definition relies either on the ciliated or the CD45^−^ signatures of tissue damage). We found that the levels of program R are significantly correlated with the viral burden (Pearson’s *r* >0.64, 0.86, *p* <10^−19^, 10^−14^ for the H1N1 and H3N2 cohorts, respectively), whereas the levels of program T are not (Pearson’s *p* > 0.05; **Figure 2D-I**). We further found that the levels of program T are significantly correlated with disease-tolerance phenotypes (Pearson’s *p* <0.05 in all cases), whereas the levels of program R are associated with disease-tolerance phenotypes only in the H1N1 cohort (*p* <0.05) but not the H3N2 cohort (Pearson’s *p* >0.2) (**Figures 2D-II, S6CD**). Thus, programs T/R and disease-tolerance/resistance mechanisms have the same signature of associations with organismal phenotypes (**Figure S5D**).

**Figure 2.**
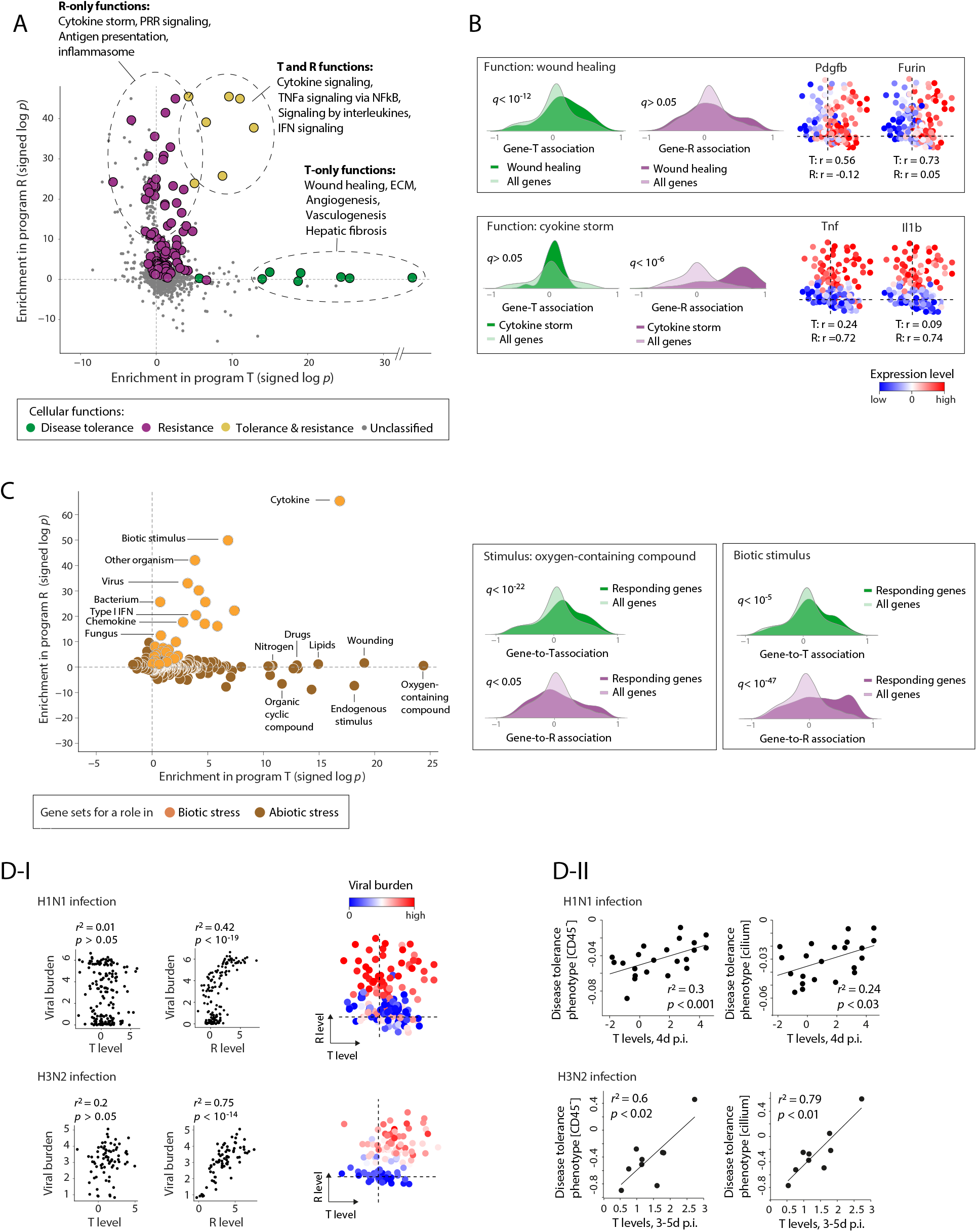
The identified T and R programs are associated with the hallmarks of disease tolerance and resistance, respectively. (**A**) Hallmark functions. For each functional gene set (a dot), indicated is the enrichment of this gene set within T-associated genes (x axis) and R-associated genes (y axis) (Wilcoxon test, signed log *p* values, positive/negative scores for enrichment within genes that are positively/negatively-associated with the program). Established disease-tolerance and resistance functions are color-coded. (**B**) Selected functions. For each function (i.e., a gene set), shown are the distributions of gene associations for genes in this gene set (dark coloring) compared to all genes (light coloring; left/right for associations with T/R), highlighting enrichments of functions in R and T genes (enrichment p-values are indicated). Right: Plots of two representative genes from each gene set (shown as in Figure 1F), highlighting associations of specific genes with programs R or T. (**C**) Hallmark response to stressors. Left: Enrichments of genes that have a known role in a certain stress response (a dot) with program T (*x* axis) and R (*y* axis). Gene sets of known roles in biotic and abiotic stress responses are color-coded. Right: selected examples, shown as in B. (**D**) Hallmark phenotypes. (**I**) Program R is linked to the viral burden. Left: viral burden against T or R levels across the H1N1-infected (top) and H3N2-infected (bottom) mice. Right: shown is the landscape of T/R levels across CC individuals (shown as in Figure 1F), with individuals (dots) colored by their measured levels of the viral burden (blue/red indicating low/high viral burden for each individual). (**II**) Program T is linked to phenotypic diversity in disease tolerance. T levels (*x* axis) versus disease-tolerance phenotypes (*y* axis) across CC strains (dots), during either H1N1 (top) or H3N2 (bottom) IAV infection. ‘Disease-tolerance phenotypes’ were calculated using the reaction norms approach (**Methods**, **Figure S6AB**). Overall, programs T and R match the hallmark functions (A,B), responses (C) and phenotypes (D) of the disease-tolerance and resistance strategies, respectively (see also **Figure S5C,D**).

Collectively, these observations suggest that programs T and R manifest the various hallmarks of disease tolerance and resistance strategies, respectively. Following these findings, the T and R programs are denoted ‘disease-tolerance’ and ‘resistance’ programs, respectively. Notably, the classical disease tolerance term points to multiple distinct mechanisms^1–4,6^. Our findings show that many of these mechanisms are associated with the same regulatory program.

### The identified programs are part of a generic molecular response in both immune and non-immune cell types

We have shown that T and R levels are related to the disease-tolerance and resistance strategies. However, the programs were observed at the bulk-tissue level and could, in principle, be due to changes in cell-type composition rather than molecular states. We therefore asked whether the T and R programs describe intracellular (cell autonomous) molecular states and in which cell types. To address these questions, we investigated the T and R programs using transcription profiles from 26 immune cell types isolated from blood^26^ (248 human individuals, either healthy or five autoimmune disease subjects), as well as primary human bronchial epithelial cells (HBECs)^27^ (120 samples, both uninfected and IAV-infected cells) (**Methods**).

We found a highly significant diversity of T and R levels in all cell types under study (**Figure S7A**). In HBECs, the dynamics of programs T and R during IAV infection is aligned with the observed dynamics in lung samples (**Figure S7B**). Thus, the T and R programs are observed in many different cell types. However, this finding relies on integration of measurements across thousands of genes and could in principle be due to different underlying genes in each cell type. To discriminate between shared and distinct responding genes of the T/R programs across cell types, we calculated, in each cell type, the association of each gene to the levels of each program across human individuals. By comparing associations of different cell types, we found that the T/R-associations are largely consistent across all human cell types under study, and strongly validate the T/R-associations in murine IAV-infected lungs (averaged associations across individuals of all human cohorts: **Figure 3A**; separate analysis of the healthy cohort and several disease cohorts: **Figure S7C,D**). For example, the associations of genes in pDCs and naive CD8 cells are highly similar and cross-validate each other (*r*=0.88, 0.77 for associations with T and R levels, respectively; **Figure 3A, right**). Given that the signatures of programs T and R are consistent across multiple isolated cell types, we conclude that both programs are part of a generic infection defense response. While it is known that specific pathways (such as interferon signaling) are shared across nearly all cell types^28^, the generality of the separation between the molecular T and R cell states has not been described.

**Figure 3.**
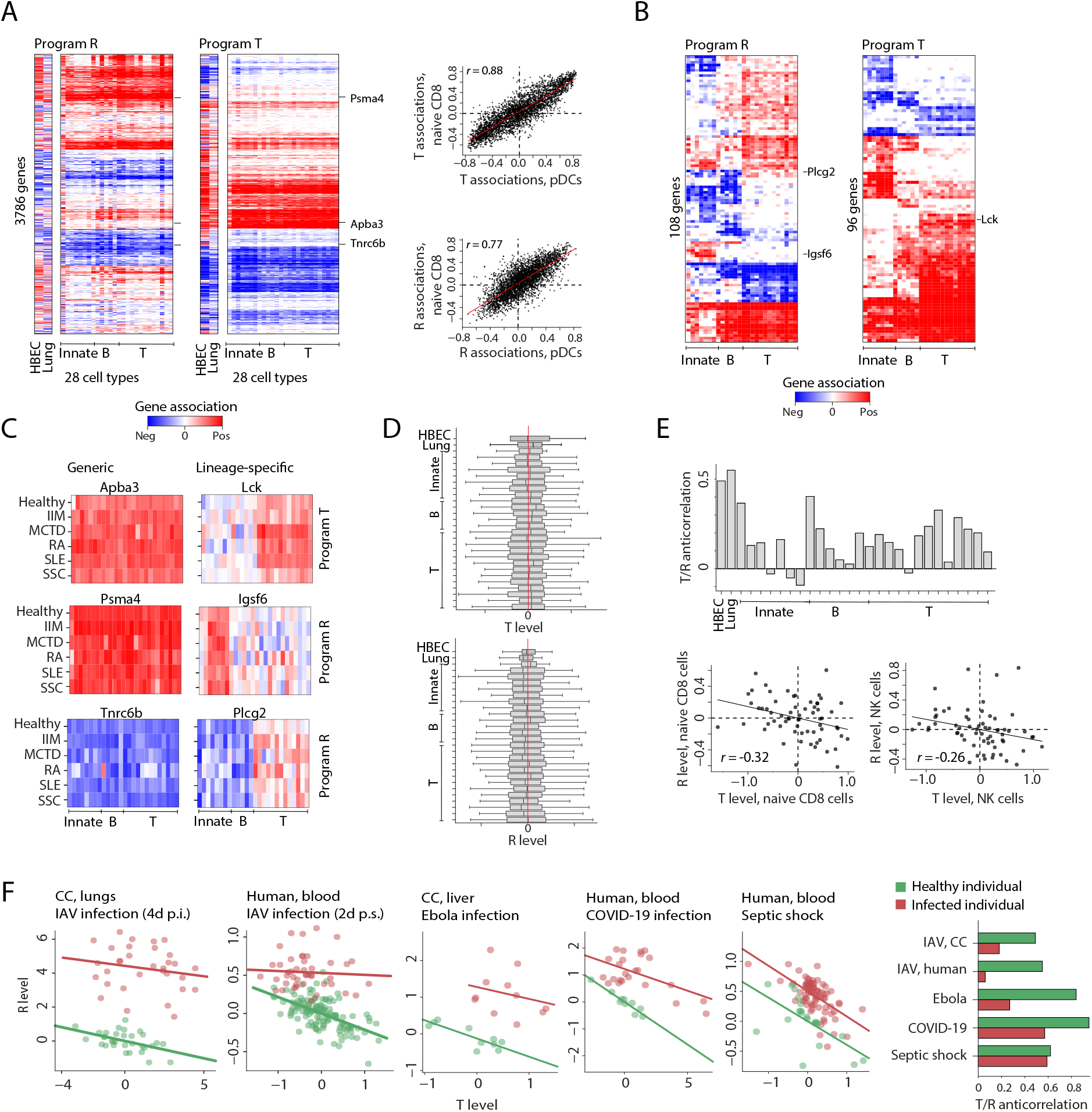
The landscape of molecular T and R states in immune and non-immune cell types. (**A**) Consistency of the T and R programs in all cell types under study. Shown are the gene-to-program associations (color-coded) for each gene (row) and each program (indicated on top), calculated based on transcription profiles in each isolated cell type (column; primary human bronchial epithelial cells [HBECs]^27^, CC lungs, and 26 human blood-derived immune cells types for which associations were averaged across six different human cohorts^26^). Right: the associations of genes (dots) with a program are consistent in two representative cell types. (**B**) A focus on genes (rows) whose T/R-associations significantly differ between cell lineages. Matrices are shown as in A. (**C**) For the indicated genes (from A,B), shown are association profiles based on data in different human cell types (columns) across six different autoimmune-disease cohorts (rows), emphasizing differences in T/R signatures between cell lineages. Left/right – generic/lineage-specific genes. (**D**) Distributions of T and R levels at the baseline state (across 63 healthy human subjects), in different cell types, demonstrating tonic activation of the programs in most healthy subjects. (**E**) T-R relations. Shown are the anticorrelations between T and R levels across healthy individuals (*y* axis, Pearson’s *r*), calculated separately using profiling of each cell type (*x* axis). Bottom: demonstration of the anticorrelation in two selected immune cell types, across healthy human subjects (dots). (**F**) The T-R balance in healthy versus infected subjects. T/R levels (*x*/*y* axes) for individuals with infectious diseases (red) and the healthy controls of each study (green). Right: summary of the T-R anticorrelation in infected (red) and healthy (green) subjects. The plots suggest a substantial T-R anticorrelation in healthy subjects that is reduced during infection (paired t-test *p* <0.02).

Notably, we did find a relatively small number of genes associated with T or R but only in specific cell types. Particularly, 5% of the genes (204 of 3786) are associated with T or R but not in all three main cell-lineages (T, B and innate immune cells; ANOVA *p* <10^−15^; **Figure 3B**). For example, these genes include Lck, which has a role in maturation of developing T cells^29^ and is indeed positively associated with program T only in T cells, and the Plcg2 gene, which is a phospholipase and is associated with program R but only in B cells and innate immune cells (**Figure 3C**). Thus, the analysis shows that the T and R responses are largely shared between cell types, and further highlights genes associated with programs T and R in a cell-lineage-specific manner.

### Organizational principles of the T and R programs

The observation that both T and R levels reflect expression states allowed us to interrogate the activation of these programs in health and disease. Interestingly, we found substantial variation in both programs that is not limited to inflammation but is also observed during normal conditions: first, in all cell types, there is a significant diversity of T and R states in healthy subjects (**Figures 3D, S7A**), and second, the diversity within healthy subjects is sufficient to demonstrate the shared gene-signature of each program across cell types (**Figure S7C**). These observations complement and extend previous studies that have demonstrated the role of specific inflammatory pathways, such as the interferon signaling, in maintenance of homeostatic balance^30,31^.

We next asked how the combination of programs T and R shapes the cell state in homeostatic conditions. In healthy subjects, we found that T and R levels are largely anticorrelated in all immune cell types (**Figure 3E**). The baseline anticorrelation also appears in human blood, lungs and liver samples in response to various pathogens (**Figure 3F**). This widespread anticorrelation is in agreement with previous genetic analyses^32^, and is also consistent with the rationale that the disease-tolerance and resistance strategies have two contradicting roles: whereas disease tolerance has a role in maintenance of homeostasis, resistance allows deviation from homeostasis to eliminate the invading pathogen. Thus, the healthy steady state is characterized by anti-correlation between the inferred molecular levels of programs T and R.

The antagonistic T-R crosstalk may seem to contradict the view that disease tolerance and resistance are two complementary strategies. We did however observe substantially reduced anticorrelation within disease patients (*p* <0.02, paired t-test; **Figure 3F**). For example, the T/R-anticorrelation is reduced during human IAV infection (Pearson’s *r* =-0.6) compared to the healthy subjects (*r* =-0.1). The same pattern (strong anticorrelation in health and a variety of T-R relations during infection) also appears in other diseases such as Ebola infection (liver samples) and SARS-CoV-2 infection (blood samples). Thus, we conclude that the antagonistic T-R relations are mostly pronounced in the healthy steady state.

### The baseline T and R states in blood MFs is associated with pathophysiology in response to infection and injury

We next used the quantitative T/R metrics to test how baseline levels of T and R are linked to future disease. Specifically, we used data from a different cohort^33^ (the BXD mouse strains^34^), which allowed us to compare between the basal T/R levels in blood-derived MFs from healthy individuals and phenotypes in these strains following injury (abiotic stimulus) and infection (biotic stimulus).

We first investigated the programs in the context of hepatic injury. Hepatic injury may lead to two different outcomes: mild injury typically leads to tissue repair (wound healing), but a repetitive injury or chronic wound may lead to fibrosis, with important roles of MFs in these processes^35^. To explore these distinct response states, we analyzed a collection of tissue-pathophysiology markers following hepatic injury across the BXD strains, including phenotypes of an aberrant wound healing (tissue damage) following a mild, transient hepatic injury, as well as fibrosis susceptibility markers following profibrotic/repetitive hepatic injury (**Table S3**). We found that a high baseline T level of MFs is associated with a beneficial state (lesser tissue damage following a transient, mild injury, *p* <10^−4^, *t*-test), but is also associated with an unfavorable state (fibrosis following a profibrotic/repetitive injury, *p* <10^−6^, *t*-test; **Figure 4A**). Thus, the baseline state of program T in blood MFs is associated with tissue pathophysiology following hepatic injury.

**Figure 4.**
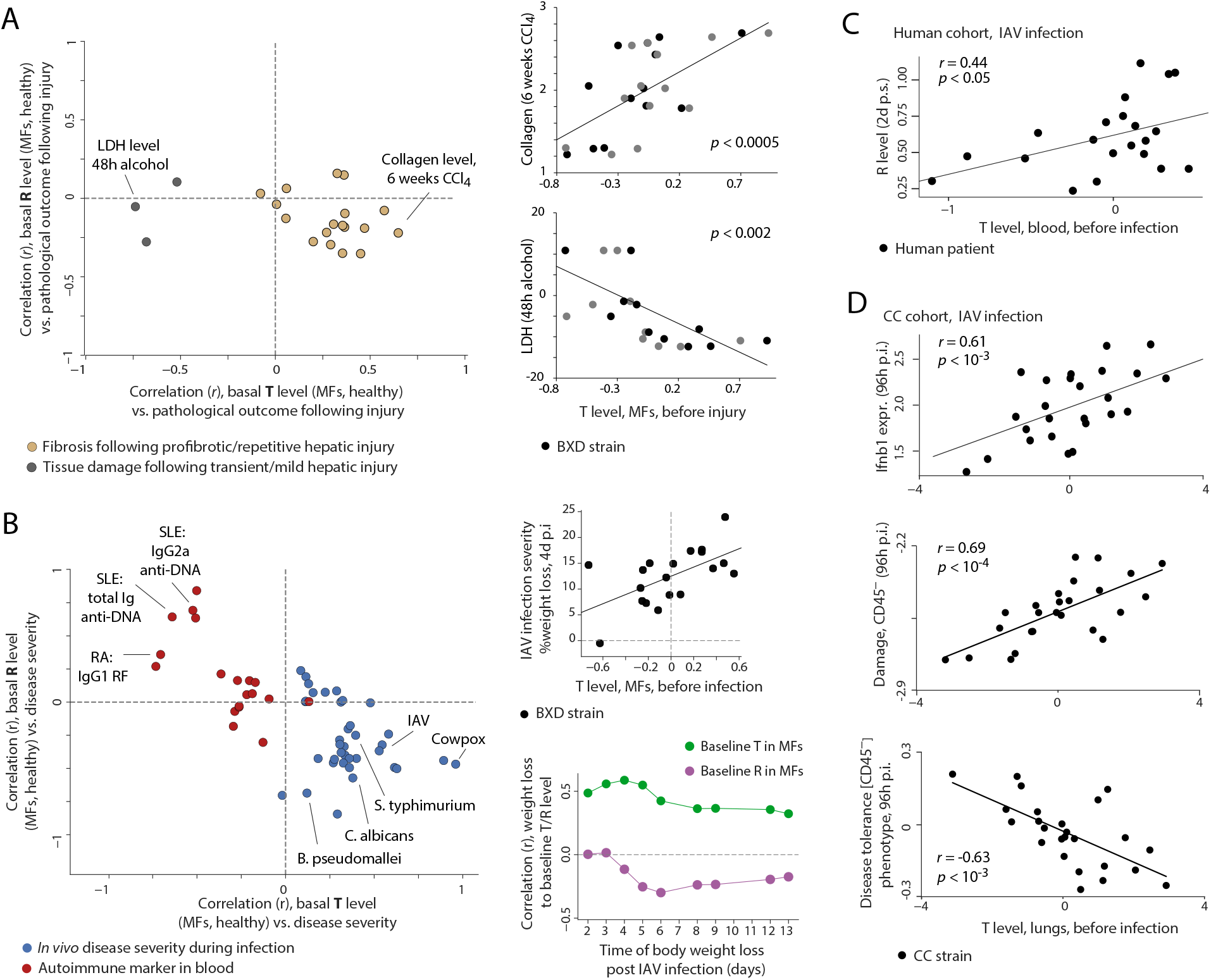
Baseline T and R states in blood MFs are linked to the *in vivo* response to stimuli. (**A**) Response to injury. Left: For various phenotypes (dots), shown are correlations between the phenotype and the baseline T (*x* axis) or R (*y* axis) levels. Correlations were calculated across the BXD strains. T and R levels were measured in resting MFs derived from the blood of healthy BXD mice. Phenotypes included are fibrosis biomarkers following profibrotic/repetitive injury (yellow) and tissue damage markers following mild/transient injury (dark gray). Right: selected examples, demonstrating baseline T levels (*x* axis) versus phenotypes (*y* axis) across mouse strains (dots), using T levels in resting (gray) and activated (black) MFs. (**B**) Response to biotic stimulus. Left: For various phenotype (dots), shown are correlations between the phenotype and the baseline T (*x* axis) or R (*y* axis) levels. Correlations were calculated across the BXD strains. T and R levels were measured in activated MFs derived from the blood of healthy BXD mice. Phenotypes included are disease severity phenotypes following infection (blue) and blood levels of autoimmune markers (red). Right: selected examples. (**C**) Phenotypic states during IAV infection in human. For human individuals (dots), shown are the relationships between T levels in blood before infection and R levels (representing disease severity) during IAV infection, at 48h post symptom onset (p.s.). (**D**) Phenotypic states during IAV infection in CC mice. Shown are the relationships between T levels in lungs before infection against disease phenotypes at 96h p.i.

We next analyzed phenotypes of *in vivo* response to various infectious diseases (viral, bacterial and fungal infections; **Table S3**). We found that higher baseline T levels in MFs were associated with a greater severity of future infections (*p* <10^−12^*, t*-test; **Figure 4B**). We confirmed the reproducibility of this finding in three independent IAV infection cohorts: using multiple time points in the BXD cohort (**Figure 4B, right**), a cohort of human patients (**Figure 4C**), and multiple disease phenotypes across the CC cohort (**Figures 4D, S5E**, focusing on Mx1-deficient CC mice to avoid confounding factors; see details in **Supp. Information 2**). Program R presents the opposite relations with the severity of illness (*p* <10^−7^, *t*-test; **Figures 4B, S5E**), consistent with the antagonistic T/R balance (**Figure 3**). Importantly, the analysis in all three cohorts indicated that disease severity is better correlated with the baseline T levels rather than the baseline R levels (**Supp. Information 2**), highlighting a link between the baseline T states in MFs and the host’s response to infection. Taken together, our findings suggest that an early ‘hit hard, hit quickly’ strategy (high resistance state against the pathogen with reduced inhibition of the resistance state by program T) is beneficial against pathogenic infections. We note that the low-T/high-R state is not always a beneficial immune state – for instance, it could be unfavorable in the context of autoimmunity (**Figure 4B**).

Overall, the analysis suggests that a dysregulated baseline T program in MFs is associated with various pathologies: very high baseline T levels in MFs are linked to severe infection and fibrosis, whereas very low baseline T levels in MFs are linked to autoimmune markers and an aberrant wound healing following a transient/mild injury. These findings suggest that an intermediate T level is beneficial as a compromise between opposing forces. In agreement, intermediate T levels are indeed highly prevalent in human healthy subjects (**Figure 3D**). Consistent with this hypothesis, tonic activation of NFkB/IFN-signaling is a known critical component of the homeostatic balance^17,31,36^.

### Arhgdia regulates the molecular T response in IAV-infected epithelial cells

To identify regulators that may induce T and R activation, we ranked possible markers of these programs. The ranking was based on consistent associations of regulators with the T or R programs across multiple cell types and contexts (**Figure 5A**, **Methods**; a complete list of markers is provided in **Table S4**). Validated IAV restriction factors, which have an established effect on the virus replication cycle^37^, indeed obtained high ranking in resistance, as expected (*p* <0.04, Wilcoxon test; **Figures 5A, S5F**). In addition, our ranking suggests a novel role for Arhgdia and Map2k2 in disease tolerance. We focused on Arhgdia, a regulator of Rho proteins, because it showed the best (positive) association with program T in the relevant experimental setting (IAV-infected murine lungs and human bronchial epithelial cells; **Figure 5A**).

**Figure 5.**
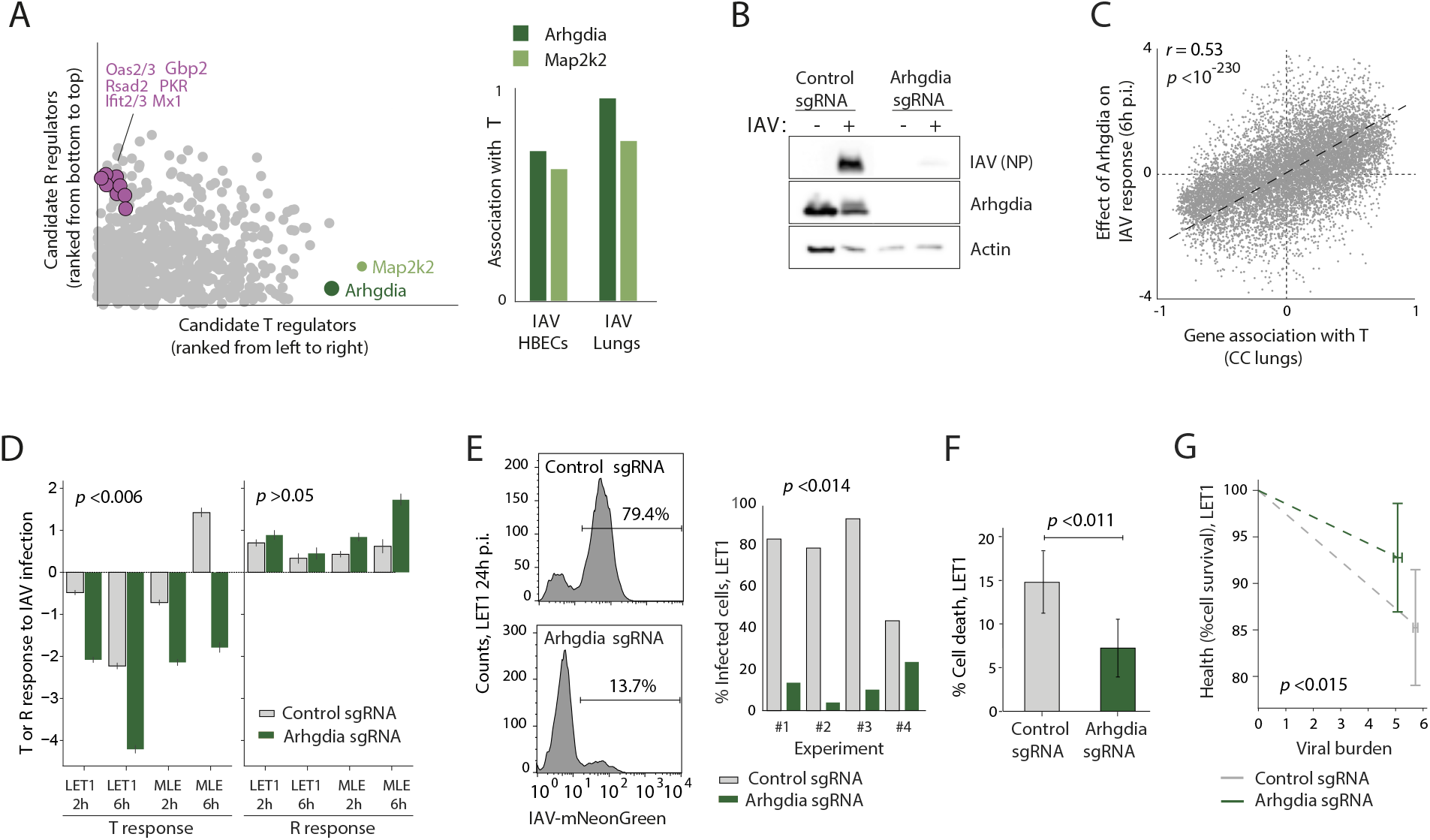
Arhgdia affects disease-tolerance responses in IAV-infected epithelial cells. (**A**) Ranking of regulator candidates by their associations with programs T and R. Validated IAV restriction factors (purple) obtained high ranking in program R; Arhgdia and Map2k2 (green) are the top-ranked genes in program T. Right: Arhgdia obtained the best T-associations in IAV-infected epithelial cells (HBECs) and CC lungs. (**B**) Immunoblot analysis of Arhgdia knockout (CRISPR-Cas9 editing) in LET1 cells and reduction in IAV infection in Arhgdia knockout cells. (**C**) For each gene (a dot), shown is its association with T levels (using lung data across CC mice; *x* axis) compared to the effect of Arhgdia on this gene (differential expression in LET1 cells: response at 6h p.i. of control versus Arhgdia-deleted cells, *y* axis). The plot shows that the effect of Arhgdia resembles the signature of program T. (**D**) Shown are T and R responses to IAV infection in Arhgdia-deleted and control cells, either MLE-12 or LET1 cells, at the indicated time points (2-6h p.i.). The plot indicates a positive role for Arhgdia in the regulation of program T (T: *p* < 0.006, R: *p* >0.05, paired t-test). Confidence intervals were calculated using bootstrapping of genes. (**E**) Representative flow cytometry data of LET1 cells infected with IAV-mNeonGreen virus (PR8 strain) for 24 hr (histograms). Bar graph (right), represents % infected cells in four independent experiments. (**F**) Shown is the percentage of dead cells, which was determined by flow cytometry using the Live/Dead assay in Arhgdia-deleted cells versus control cells in 4 independent experiments (24h p.i.). (**G**) Comparison between the ‘disease-tolerance phenotype’ in Arhgdia-deleted LET1 cells versus control LET1 cells at 24h p.i.. The disease-tolerance phenotype is defined as the slope of cell-survival against the viral burden. The plot shows a higher disease-tolerance phenotype for Arhgdia-deletion compared to control cells (F-test *p* <0.02). Plots E-G indicate that the effect of a baseline Arhgdia deletion is beneficial (leading to reduced infection severity and improved ability to tolerate infection), consistent with the predicted effects of baseline T levels in Figure 4. In all plots, MOI = 5 of PR8 virus, titrated on MDCK cells.

To directly validate the power of our model, we functionally assessed the role of Arhgdia in the regulation of program T. We deleted Arhgdia in mouse epithelial cells (LET1 and MLE cells) by CRISPR-Cas9 editing, and then tested the effect of this deletion on the host response to infection. Control (non-targeting sgRNA) and Arhgdia-deleted (Arhgdia-targeted sgRNA) cells, with and without IAV infection, were analyzed. We confirmed that LET1 and MLE-12 cells express Arhgdia and that upon knock out its expression is eliminated (**Figures 5B**, **S8A**). The effect of Arhgdia on programs T and R was evaluated (using transcriptomes) at early stages of infection (2-6h p.i., **Figure S8B**) in an attempt to avoid effects of cellular damage on the overall gene-expression state. To assess the effect of Arhgdia on disease severity, we measured cell damage (using the percentages of cell death) and viral burden (quantified in infected cells by the expression of the mNeonGreen fluorescent marker, expressed by a recombinant PR8 virus, named ‘PR8-mNeonGreen’); both recorded by FACS analysis of IAV-infected LET1 cells at 24h p.i (**Methods**).

Several lines of evidence support a positive effect of Arhgdia on the T state (rather than the R state). First, Arhgdia’s responding genes are enriched with T-associated functions such as wound healing, ECM and hepatic fibrosis (hyper-geometric test *p* <10^−11^, 10^−6^, 10^−28^, respectively). Second, the effect of Arhgdia on the expression of genes strongly resembled the associations of genes with program T (e.g., Pearson’s *r* =0.53, *p* <10^−230^; **Figure 5C**) rather than program R (*r* =- 0.1, **Figure S8C**). Indeed, deletion of Arhgdia led to a significant, greater-than-normal downregulation of T levels, but the effect on R levels was weak (*p* <0.006 for T, *p* >0.05 for R, paired t-test; **Figure 5D**). Third, Arhgdia causes the same effect on disease phenotypes as the effect of program T. Particularly, Arhgdia-deletion was beneficial, leading to reduced viral burden (*p* <0.014, **Figure 5B** [Immunoblot analysis] and **Figure 5E, S8D,E** [flow cytometry analysis]) and reduced cell death (*p* <0.011, **Figures 5F**), consistent with the observation that a lower early T level is linked to a less severe *in vivo* infection (cf. **Figure 4B-D, S5E**). Finally, we calculated the ‘disease tolerance phenotype’, defined at the cellular level as the slope of cell survival against viral burden. We found that the presence of Arhgdia is linked to a lower disease-tolerance phenotype (F test *p* <0.015; **Figure 5G**), consistent with the observation that higher baseline T levels are linked to a lower ability to tolerate future *in vivo* infections (**Figure 4D**). These observations provide additional support for our conclusion that program T (and Arhgdia as one of its regulators) is associated with alterations in the phenotype of disease tolerance during IAV infection.

## CONCLUDING REMARKS

In this study we explored gene signatures that are linked to the two main defense strategies: disease tolerance and resistance. We identified a gene program for the disease tolerance strategy (T) that is separable from the gene program of the resistance strategy (R). We further provided refinement for the current gene signature of resistance by allowing its precise definition that is uncoupled from the gene signature of disease tolerance. These definitions allowed us to study, for the first time, the relations between the two defense strategies at the molecular level. We found that both T and R states are robustly detected at the gene-expression level in multiple cell types, including non-immune, innate and adaptive immune cell types. The analysis revealed generic signatures for the identified T and R states across the various cell types (both mouse and human cells), allowing us to develop personalized and quantitative metrics for the molecular level of each program. The two programs explain a roughly equal percentage of the molecular and phenotypic variation, consistent with previous studies that have shown a roughly equal number of disease-tolerance and resistance mutations that influence the progression of infections^38^.

We found a tight link between the baseline T levels in MFs (before stimulus) and the *in vivo* pathological response following stimulations. While excessive baseline T activity in MFs is correlated with elevated pathology in case of chronic injury and infections, the baseline MF state of poor T levels is correlated with elevated pathology in response to transient injury. We speculate that the homeostatic intermediate levels of program T may benefit from the compromise between opposing forces. In line with this, there is a high prevalence of the intermediate T level in healthy subjects. Further work is needed to characterize the associations of programs T and R with additional disease outcomes and therapeutic responses. For example, one additional physiological condition in which the T program could be very important is pregnancy: it has been earlier shown that a strong pro-inflammatory phenotype may lead to a spontaneous miscarriage^39,40^, and a counterbalancing T program is potentially needed for a successful pregnancy.

Our data suggest that a high-T/low-R state before infection is associated with a greater severity of future infections, implying a central role of the baseline T-R balance in disease progression. These findings further suggest a ‘hit hard, hit quickly’ model of the host defense against pathogens: a greater severity of infection could be a consequence of an early inhibition of resistance by disease-tolerance functions. It has been long argued whether death resulting from dysregulation of inflammation in sepsis is due to a poor initial immune response followed by high pathogen burden and secondary hyper-inflammation or, alternatively, is due to a disproportionately strong initial immune response against a low pathogen burden that leads to inadequate systemic responses. Our findings argue for the former, in line with studies of bacterial sepsis^41^ and respiratory infections^42–44^, and consistent with the efficacy of approaches that boost the baseline/early resistance such as vaccines^45,46^.

We also demonstrate the robustness of our framework through experimental validations of Arhgdia as a key regulator of the cellular T program. Arhgdia was co-expressed with program T in all cell types and under various conditions. Arhgdia is known as a regulator of Rho proteins and its activity improves survival of stem cells^47^ and kidney functions^48^. However, its more general role in control of disease tolerance has not been reported. Consistent with our predictions, we identified a novel role for Arhgdia in regulation of the cellular T state. The approach presented here, particularly the specific scores (and markers) of T and R levels, is a generally applicable framework. The approach can therefore be used to identify novel regulators and therapies that specifically target the balance between disease tolerance and resistance states at the molecular level.

## METHODS

### Longitudinal IAV infection data across CC mice

#### Mice

The CC cohort includes female mice aged 7–10 weeks from the Tel Aviv University (TAU) collection of Collaborative Cross recombinant inbred mice^49^, as well as the C57BL/6J strain from Envigo, Israel. The mice were raised at the Animal Facility at the Sackler Faculty of Medicine of TAU. All experimental procedures were approved by the Institutional Animal Care and Use Committee (IACUC) of TAU (approval number 04-14-049) and adhere to the Israeli guidelines and the US NIH animal care and use protocols. Mice were held in individually ventilated cages and housed on hardwood chip bedding under a 12h light/dark cycle, humidity-controlled and temperature-controlled conditions. Mice were given tap water and standard rodent chow diet ad libitum from their weaning day until the end of the experiment.

#### In vivo IAV infection

Mouse-adapted H1N1 influenza A/PR/8/34 virus was grown in allantoic fluid of 10-day-old embryonated chicken eggs at 37°C for 72 h. Allantoic fluid was harvested and viral titres were determined by standard plaque assay in Madin–Darby canine kidney (MDCK). All mice were first anesthetized (intraperitoneally) with 7 mg ml–1 ketamine and 1.4 mg ml– 1 xylazine at 0.1 ml per 10g body weight. Next, animals were infected intranasally with PR8 (4.8 × 10^3^ pfu in 40 µl PBS). The data consist of one mouse from each strain and each time point. Over the time course for 33 CC strains, data were missing for only a few individuals. For additional six strains, only one or two individuals were evaluated at late time points of infection (detailed in **Table S1**).

#### Organismal phenotypes

Whole-body weight and lungs’ functions were measured daily. Respiratory parameters were measured using an unrestrained whole-body plethysmography system (DSI, MN, USA). A non-anesthetized mouse was placed in a plethysmography chamber and acclimated for 20 min. Respiratory parameters were then collected for an additional 5 min, and average values were used for the enhanced pause (Penh) parameter to assess airway dysfunction as previously described^50^.

#### Measurement of gene expression

Lung tissues were collected into RNAlater (Qiagen). Lysis was performed with QIAzol (Qiagen). RNA isolation was performed according to miRNeasy kit protocol (Qiagen). mRNA quality was checked using the Agilent 2100 Bioanalyzer according to the manufacturer’s instructions. All RNA Integrity Numbers (RINs) were higher than 8. cDNA libraries were prepared from 2 µg of isolated RNA using the SENSE mRNA-Seq Library Prep Kit V2 for Illumina (Lexogen). Each sample had its own index primer. DNA size and quality were checked using the Agilent 2100 Bioanalyzer. Libraries were quantified using the Qubit DNA HS Assay kit (Invitrogen). The amplified libraries were pooled at a total concentration of 2 nM and sequenced using the Illumina HiSeq platform at the Technion Genome Center (Israel). This dataset is deposited in the GEO database (GSE174253).

#### Pre-processing of gene expression

We applied a joint alignment of sequencing reads for both the mouse genome and the influenza virus genome using the Bowtie2 software and then applied a joint quantification of both the virus and mouse transcripts. For this quantification, we applied an FPKM-normalization using the RSEM software, and then ln-transformed the data. Additional standardization of expression levels (centered and divided by standard deviation) was applied across genes (sample-level) and then across samples (gene-level) based on the data collected before infection. Unless stated otherwise, the ‘expression levels’ refer to these standardized log-transformed gene expression levels.

#### Expression-derived disease severity phenotypes

Disease severity was quantified by analyses of viral burden, %weight loss, breathing dysfunction, *Ifnb1* and *Ccl2* expression, the quantity of immune cells, and tissue damage. The ‘viral burden’ phenotype was defined as the averaged expression level of viral transcripts NC_002016, NC_002017, NC_002018, NC_002019, NC_002020, NC_002021, NC_002022, and NC_002023; values were *ln*-transformed but were not standardized. ‘Weight loss’ is reported as the percentage of reduction in the whole-body weight compared to the weight immediately prior to infection. ‘Breathing dysfunction’ is defined by the Penh metric^50^. The expression levels of *Ifnb1* and *Ccl2* were obtained from the transcriptome data (*ln*-transformed, with standardization). The mean expression levels of CD45^+^-specific genes were used as a signature for the quantity of immune cells, and the negation of mean expression levels of genes specific to CD45^−^ cells was used as a signature for tissue damage. To define the sets of CD45^+^-specific and CD45^−^-specific genes, we first used Seuret v3^51^ to annotate each single cell from the Tabula Muris dataset^52^, and then selected lung cells that did or did not express CD45. The CD45^+^-specific genes were defined as the genes at the top 25% of average expression in the CD45^+^ cells and at the bottom 25% of average expression across the CD45^−^ cells. Similarly, the CD45^−^-specific genes were defined as a set of genes at the top 25% of average expression across the CD45^−^ cells and at the bottom 25% of average expression across the CD45^+^ cells. Tissue damage was defined based on two metrics: the average reduction in the expression of CD45^−^-specific genes and the average reduction in the expression of cilium genes. We further used a signature of ‘cytokine storm’, which was defined as the average expression of the set of cytokines typically dysregulated during cytokine storm syndrome^22^. Finally, the Mx1 genetics of each strain was determined as previously described^20^.

#### Assessment of inherited variation

To calculate the heritability measure (variation due to genetic differences) for each phenotype, we quantified two mice per strain on day 4 post IAV infection (12 CC strains). Measurements of gene expression were collected and pre-processes as previously described^53^. As the main longitudinal CC dataset was generated using a different technology (see above), this limited 4-days dataset was only used for the assessment of inherited variation. This dataset is deposited in the GEO database (GSE184278). A p-value of heritability was calculated using comparison to a null distribution of the heritability score, which was calculated for each phenotype using permutation of individuals between strains.

#### Assessment of the ‘disease-tolerance phenotype’

The disease tolerance phenotype is generally defined as the level of health relative to the pathogen burden^25^. At the broad sense, this phenotype is calculated based on the entire physiological path in the health space^14^. However, most datasets do not contain the entire course of infection but only one or a few time points (e.g., in our case, the data includes the incubation and acute phases but not the outcome phase [see **Figure S1B**], a limitation imposed by ethics considerations) and therefore it is impossible to analyze the entire health continuum.

An alternative approach, commonly used in the relevant literature, relies on reaction norms^6,14,25^. Reaction norms plot the level of health for an individual at each pathogen burden. The “disease tolerance phenotype” is defined as the slope of the health-to-pathogen regression in this plot, such that a shallower slope indicates a better ability to tolerate the pathogen (see examples in **Figure S6AB**). The analysis of disease-tolerance phenotypes compares the slope between groups of individuals. To calculate the slope for a given group, the group must contain a wide spectrum of pathogen burden and health levels. To address this, previous studies commonly used multiple genetically distinct individuals in the same time point^32^, or alternatively, multiple individuals from the same strain in several time points^23,54^ (it is also common to use multiple individuals from the same strain and the same time point, but only when the pathogen burden is constant^55^).

In this study, the ‘disease tolerance phenotype’ was calculated by the slope of each CC strain using a group of multiple infected individuals from the same strain (see examples in **Figure S6A,B**). We used the viral mRNA level as the measure of pathogen burden (as in **Figure 1B**) and used the negation of the tissue-damage signature as the measure of health. The tissue damage signature relies on either CD45^−^ cells or ciliated cells (see definition above and **Figures 1B, S1A**). Only one slope (H1N1-infected 6020A strain) deviated by >2 interquartile range (IQR) and was therefore excluded.

### Construction and metrics of programs T and R

We aimed to construct a model representing the co-regulation of genes during the course of IAV infection. The analysis consists of two steps. First, construction of a map, which allowed us to define the T and R programs, and second, calculation of tissue-immune states based on programs T and R.

#### Construction of a map

Our goals were twofold: first, to integrate multiple time points and strains into one representative model, and second, to construct a model relevant to the severity of IAV infection. The map was constructed in three steps. In step 1, we selected genes that have high relevance to disease severity and that provide a uniform coverage of behaviors across time points and strains as follows: (*i*) We calculated the Pearson’s correlation between the weight loss phenotype at 96h p.i. and the expression of each gene across mouse samples (with a separate calculation at each time point). The output is a matrix of correlations for each gene at each time point. (*ii*) We selected genes with relevance to disease severity as those with absolute correlation with the weight loss higher than 0.15 and with the same direction of correlation in two consecutive time points. Of 24,500 genes, 5075 genes were retained in the correlation matrix. (*iii*) We applied hierarchical clustering on the correlation matrix resulting in 15 main clusters of genes. (*iv*) For each cluster and each time point, we identified a ‘core’ group of co-regulated genes. Each core is a set of 30 genes whose average correlation with the remaining genes in the same cluster is the highest. There were 75 cores, one for each cluster at each time point. (*v*) For each individual mouse, we calculated the average expression across the 30 genes in each core at the relevant time point, resulting in a vector of 75 values for each individual mouse, which represents the main co-regulation patterns across all time points.

In the second step, we constructed a core-centered representation in which each gene was represented by its interrelations with the 75 core signatures. Specifically, we calculated the correlation between each core and each of the 5075 genes across individuals (using gene expression at 96h p.i.).

In the third step, we constructed a map based on this core-centered representation of genes. Using the core-centered representation of genes as input, we trained an autoencoder neural network with a two-dimensional bottleneck layer and a sigmoid activation function. We focused on a two-dimensional map, since the third dimension has a limited contribution to the explained variation (**Figure S1F**). We used the bottleneck layer as the two-dimensional representation, referred to here as the ‘map’. The map construction was performed in two steps: first, the encoder was trained using data from the 5075 selected genes, and then the genes that were filtered out in step 1 were projected onto the map using the trained network. Of note, the entire construction was applied without pre-standardization of expression levels. To avoid a bias toward early time points (in which the selected time intervals were shorter), the data at 3h p.i. was omitted from the construction of the map.

#### Definition of the main programs and their quantitative scores

The two-dimensional map of the host response consists of two axes that are referred to as “axis T” and “axis R”. We decomposed the observed gradient of each individual (**Figure 1D-II**) into two gradients along these axes. Decomposition was performed through a deconvolution-based approach by solving the following regression model: *g_ij_* = *T_i_x_j_* + *R_i_y_j_* + *b_i_* (Eq. 1), where *g_ij_*, *x_j_*, and *y_j_* are given as input and the *T_i_* and *R_i_* values are the output. Specifically, *g_ij_* is the (standardized) expression level of gene *j* in sample *i*, and *x_j_* and *y_j_* are the input positions of gene *j* in the map. The values of *T_i_* and *R_i_* are the output “levels” of the gradients along the T and R axes in sample *i*, respectively (*b_i_* is a sample-specific constant).

Given that expression levels are largely orchestrated along each of the axes, we reasoned that each axis represents a certain program of regulation. In accordance, the T and R axes are referred to as *programs* and the inferred levels of gradients along the T and R axes (i.e., the *T_i_*, *R_i_* values) are referred to as the *levels* of programs T and R (in short, the “T level” and “R level”), respectively. Given the observed increase in T and R levels during infection, positive levels are referred to as ‘activated’ programs, negative levels are referred to as ‘inactivated’ programs, and a zero-level of a program corresponds to an intermediate level of activation. The T and R programs/levels are also referred to as ‘disease tolerance’ and ‘resistance’ programs/levels, respectively.

### Additional datasets

We tested the generality of the model using several gene expression studies. In each dataset, pre-processing was applied as described in its original publication. For each dataset, gene expression values were standartized using the ‘healthy’ samples of the dataset, and then T and R levels were calculated for each sample. The following datasets were evaluated:

1. Isolated cells from 26 cell types derived from blood samples across 63 healthy subjects, 60 SLE patients, 45 SSc patients, 19 RA patients, 42 IIM patients and 19 MCTD patients^26^.
2. *In vitro* response to PR8 virus stimulation of primary human bronchial epithelial cells in ten time points^27^ (GEO accession GSE19392). In addition, this data contains additional stimulations (e.g., interferon, viral RNA) that were used for the calculation of epithelial gene-to-program associations (a total of 120 samples).
3. A human IAV-infection cohort, consisting of 21 individuals (12 females, 9 males), infected with IAV during the 2009-2010 influenza season^56^. We analyzed, for each individual, blood transcription profiles that were taken before the influenza season and at 0, 2, 4, 6 and 21 days after symptom onset. As clinical phenotypes are not available, we used the R level at 2 days after symptom onset as the marker of disease severity (GSE68310).
4. Lung samples of the C57BL/6J mouse strain at 10 time points following *in vivo* IAV infection^53^ (GEO accession GSE49934).
5. Skin samples of the ICR mouse strain at 8 time points (0-192 h) post injury^57^.
6. Microarray data for LPS-treated and control primary macrophages of BXD strains^33^ (GEO accession GSE38705). Phenotypes across these strains are detailed in **Table S3.** IAV infection severity was measured as the percentage of body weight loss (data from Ref. ^58^).
7. Transcriptional profiles from liver across CC mice, including Ebola-infected mice and controls^59^ (GEO accession GSE130629).
8. A human cohort of whole blood samples from children with septic shock (*n*= 82) and healthy controls (*n*=21). Samples taken within 24 hours of admission to the pediatric intensive care unit^60^ (GEO accession GSE26378).
9. A human cohort of whole blood samples from patients with community-acquired pneumonia (CAP) by SARS-CoV-2 within the first 24 h of hospital admission^61^.
10. A mouse cohort of lung samples from IAV (H3N2 strain)-infected CC mice (9 MX1-deficient strains, 3 mice per strains, profiled at 3 and 5 days post infection)^24^ (GEO accession GSE136748).

### Functional organization of the map

Functional analysis proceeded in two steps. First, we calculated the correlation of each gene to the levels of each program (a gene-to-program association score; Table S2). Then, for each gene set and each program, we calculated the bias of the associations of genes within the gene sets compared to the remaining genes (a Wilcoxon rank-sum test). The resulting ‘functional enrichment’ score is defined as the log of the Wilcoxon p-value (FDR-corrected), signed by the direction of bias, such that positive and negative signs indicate correlations that are higher and lower than the expected distribution. This functional analysis was applied systematically across several public repositories of gene expression datasets. For clarity, the gene sets were organized into four collections of distinct biological interpretations.

1. Gene sets of biological functions and processes. We used the entire collection of gene sets from MSigDB’s ‘Hallmark’ and ‘Reactome’ repositories (*n*= 1568) (**Figure 2A,B**). We classified the functions as either disease-tolerance or resistance functions.
2. Sets of genes that have a role in response to stress. We used all gene sets from the GO database with annotations that included the term ‘response to’ (356 gene sets; **Figure 2C**). We note that these sets are not limited to transcriptional response but in fact include any protein that have a known role in the cellular stress response.
3. Gene sets related to NFkB and interferon signaling, collected manually from MSigDB and the Ingenuity knowledge base (32 gene sets, **Figure S5A**).
4. Positive and negative regulators of various biological functions. Included were genesets from the GO repositories with annotations that start with either ‘positive regulation of’ or ‘negative regulation of’. Overall, we used *n*=625 pairs of gene sets, where each pair includes one geneset of positive regulators and one gene set of negative regulators for the same function (**Figure S5B**). The analysis excluded genes that encoded factors that were dual positive and negative regulators of the same function.

### Ranking of disease-tolerance and resistance regulators

We used an unbiased approach to rank candidate regulators. The ranking was performed for each program (T and R) separately. We first applied two selection criteria: (1) Whether the gene is tightly associated with the program (absolute *r* >0.8) in at least one cell type using at least one cohort (either healthy individuals or one of five autoimmune-disease cohorts : SLE, RA, SSc, IIM,MCTD; data from Ref. ^26^). (2) Whether the gene is annotated as a regulator, using annotations of “signaling molecules” “receptor”, “transcription factor” and “transcription regulator” in the Ingenuity Knowledge Base. Next, we ranked genes by their average associations to either T or R across datasets (i.e., averaging all association profiles in **Figures 3A** and **S7D**).

### Analysis of Arhgdia

#### Cell lines

Immortalized mouse Lung Epithelial Type I (LET1) (BEI Resources, NIAID, NIH, NR-42941), Madin–Darby canine kidney (MDCK), 293FT and mouse lung type II epithelial (MLE-12) cell lines were maintained in Dulbecco’s modified Eagle’s medium (DMEM, high glucose) supplemented with 10% fetal bovine serum (FBS), 2mM L-glutamine, 1% penicillin/streptomycin (all from Biological Industries, Kibbutz Beit-Haemek, Israel). All cells were grown at 37°C in a humidified atmosphere containing 5% CO_2_.

#### Viruses

Influenza A virus strain A/PR/8/34 (H1N1) and recombinant PR8-mNeonGreen (a clone named “PB2/PA PTV1 mNeonGreen”, which harbors two copies of the gene of the mNeonGreen fluorescence marker, fused to the PB2 and PA polymerase ORFs, via a porcine teschovirus-1 2A peptide sequence), were propagated by allantoic inoculation of 10-days-old chicken embryonated eggs (Kimron Veterinary Institute) and stored at −80°C. Viral titers were determined using MDCK cells and expressed as median tissue culture infective dose (TCID_50_)^62^. For infection experiments, 24-well plates were seeded with LET1 (50000 cells/well) or MLE-12 (100000 cells/well) cells. After 24h, cells were washed with PBS and infected with IAV (MOI=5), in a serum-free medium for 1hr with gentle rocking. Following infection, the inoculum was removed and cells were incubated in OptiMEM I reduced medium (Thermo Fisher Scientific) for the indicated time. All experiments were carried out in triplicate. Flow cytometry data were acquired from 10,000 gated-events using S100EXi flow cytometer (Stratedigm) and analyzed using FlowJo version v10 software (FlowJo, LLC).

#### CRISPR/Cas9 system

Lenti vectors harboring the CRISPR/Cas9 system for Arhgdia targeting were produced by co-transfection of 293FT cells with psPAX2 (Addgene, #12260), pMD2.G (Addgene, #12259), and LentiCRISPRv2 (Addgene, #52961) plasmids, using Lipofectamine 3000 (Thermo Fisher Scientific). Single-guide RNA (sgRNA) included sg-Arhgdia-#1 (5’-CACCGTGAGTTCCTGACACCCATGG-3’), and sg-Arhgdia-#2 (5’-CACCGTGAGTTCCTGACACCCATGG-3’), targeting the murine Arhgdia gene, or a non-targeting sgRNA (’control’, 5’-CACCGACGGAGGCTAAGCGTCGCAA-3’). For simplicity, sg-Arhgdia-#1 is referred to as ‘Arhgdia sgRNA’ and the non-target is referred to as ‘control sgRNA’. After 48 hr, supernatants containing virus–like particles (VLPs) were collected, filtered (0.45 μm), and used to transduce LET1 and MLE-12 cells in the presence of polybrene (8μg/ml). After 24h, cells were placed under puromycin (2 μg/ml; Sigma #P8833) selection. LET1 resistant cells were seeded into a 96-well dish (1 cell/well) for the expansion of single clones. Transduced LET1 colonies and transduced pooled MLE-12 cells were selected and verified for the loss of Arhgdia expression by immunoblotting.

#### Immunoblot analysis

Immunoblot analysis was performed with the following primary antibodies and dilutions: anti-RhoGDI (Santa Cruz Biotechnology, #sc-373723, 1:1000), anti-β-actin clone C4 (MP Biomedicals, #0869100, 1:10000), anti-Influenza A nucleoprotein (NovusBio, #NBP2-16965, 1:1000). Secondary antibodies and dilutions were Goat anti-mouse horseradish peroxidase-conjugated (Jackson ImmunoResearch, #115-035-062, 1:10000), Goat anti-Rabbit horseradish peroxidase-conjugated (Jackson ImmunoResearch, #111-035-045, 1:10000).

#### Cell viability

Cell viability was determined with Live/Dead fixable red stain kit (Thermo Fisher, #L34972).

#### RNA profiling

Library construction, sequencing and pre-processing of data were performed as described above, with the following differences: Total RNA was extracted from cells using the Direct-zol RNA Miniprep kit (ZYMO Research). RNA samples (0.5 µg); sequencing was performed using the Illumina NextSeq 550 platform (Technion Genome Center, Israel); and Alignment of IAV and mouse (GRCm39.104) chimeric genome performed using STAR 2.7.9 aligner. Data was deposited in the GEO database (GSE193160).

## Supplementary Information 1: The relevance of programs T and R in lungs

We used several tests to assess the relevance of programs T and R in lungs of CC mice throughout IAV infection. First, we ask whether the observed diversity of T and R levels is a biological effect. Comparison of the distribution of shuffling-based T and R levels to the original T and R levels show that the T and R levels are highly significant (*p* < 10^−200^, t-test; **Figure S3A**; reshuffling-based levels were calculated based on a map with reshuffled gene positions). These observations imply that the observed diversity of T and R levels is not dominated by noise but rather a biological effect. To further support this notion, we assessed the contribution of genetics to the diversity of T/R levels. Specifically, we used a limited set of 12 CC strains for which two mice were collected from each strain on day 4 post infection. We found that the variation in T and R levels that is explained by genetic differences is highly significant (heritability *p* <10^−4^, 10^−12^ for T and R, respectively; **Figure S3B**), indicating that the T/R variation is partially inherited. Taken together, we concluded that the observed diversity of T and R levels is a biological effect.

Next, we assessed the contribution of each program to the overall gene-expression diversity. Particularly, we calculated the percentage of inter-individual variation in each gene that is explained by the T and R levels. This analysis shows that both T and R capture a large percentage of the gene-level variation during infection (**Figure S3C**). For example, for 50% of the genes, T level explained more than 10% of the inter-strain variation at 4 days p.i., and R levels explained more than 20% of their inter-strain variation at 4 days p.i. These findings indicated that both programs are of biological importance.

The analysis of explained variation allowed us to test the hypothesis that programs T and R are decoupled. For each gene, we compared two models: (i) a model in which the linear combination of T and R programs is used to explain gene-level variation, and (ii) a model in which only one program (R or T) is used to explain gene-level variation. We found that the combination of T and R outperformed the models of only one program (t-test *p* <10^−100^, **Figure S3C, D**). Thus, the two programs are decoupled and each of the programs contributes to the global state. Of note, decoupling does not imply complete independence and it is possible that the two programs are interrelated (we demonstrate a cross-talk between the two programs in **Figure 3E,F**).

## Supplementary Information 2: Using independent cohorts to confirm the relations between baseline T levels before infection and the late severity of IAV infection

Here we used three independent cohorts to confirm that T levels in the baseline state (before infection) are linked to late severity of IAV infection: human blood samples^56^, mouse lung samples (CC mice, this study), and mouse blood-derived MFs (BXD mice^34^). As shown below, in all cohorts, disease severity was better correlated with baseline T levels than baseline R levels.

### Human blood

In the human cohort, expression measurements at the baseline state were taken at least a few weeks before the occurrence of IAV infection^56^. Consistent with the two other cohorts, the baseline T levels were negatively correlated with the severity of infection (**Figure 4C**). There was a better correlation of disease severity with the baseline level of T (*r* =0.44) compared to the correlation with the baseline level of R (*r* =0.1), in agreement with the observations across the BXD and CC cohorts.

### CC mice

We compared T and R levels before infection to the severity of IAV infection across the CC lines. Specifically, we focused on the wide inter-individual variation within the group of Mx1-deficient CC strains (26 of 33 strains), which is not explained by previously described genetic differences (**Figure S1D, Table S1**). Consistent with our findings in the BXD and human cohorts, we found that disease severity at the peak of symptoms (96h p.i.) was associated with T and R levels before infection: A higher activation of the R program before infection was linked to a lower disease severity, whereas a higher activation of the T program before infection was linked to a higher disease severity (similar results using all phenotypes; **Figure 4D, S5E**). The state in early infection (24h p.i.) shows the same relations with late disease severity (**Figure S5E**). Disease severity was better correlated with baseline T than baseline R (paired t-test *p* <10^−6^ across phenotypes, **Figure S5E**), emphasizing the central effect of the early T state on illness.

These observed relationships are particularly impressive when considering that each time point was profiled using a different group of individuals (one from each strain), and therefore we cannot expect to explain more than the inherited variation (see estimation for the inherited variation in **Figure S1C**). Indeed, the explained inherited variation was systematically larger than the explained total phenotypic variation, ranging from 26% to 100% explained inherited variation across phenotypes (**Table S5**). For instance, whereas the baseline T levels explained 37% of the total variation in the late expression of type 1 interferon (**Figure 4D, top**), it explained a much higher percentage (100%) of the inherited variation in this phenotype.

### BXD lines

We compared T and R levels before infection to the severity of IAV infection across the BXD lines. For each BXD line, we used (i) the inferred T and R levels before infection, calculated using transcriptomes of blood-derived primary macrophages (data from Ref.^33^), and (ii) the severity of *in vivo* IAV infection, measured as the percentage of body weight loss, 9 different time points (data from Ref. ^58^). We found that the baseline T levels have negative correlation with the severity of IAV infection, consistent with our findings in the human and CC cohorts. Weight loss was better correlated with baseline T levels than baseline R levels (paired t-test *p* <10^−3^ across time points, **Figure 4B**), emphasizing the central effect of the early T state on illness. Similar results were obtained using LPS-activated MFs (data not shown).

## Supplementary Figures and Tables Legends

**Supplementary Figure 1.**
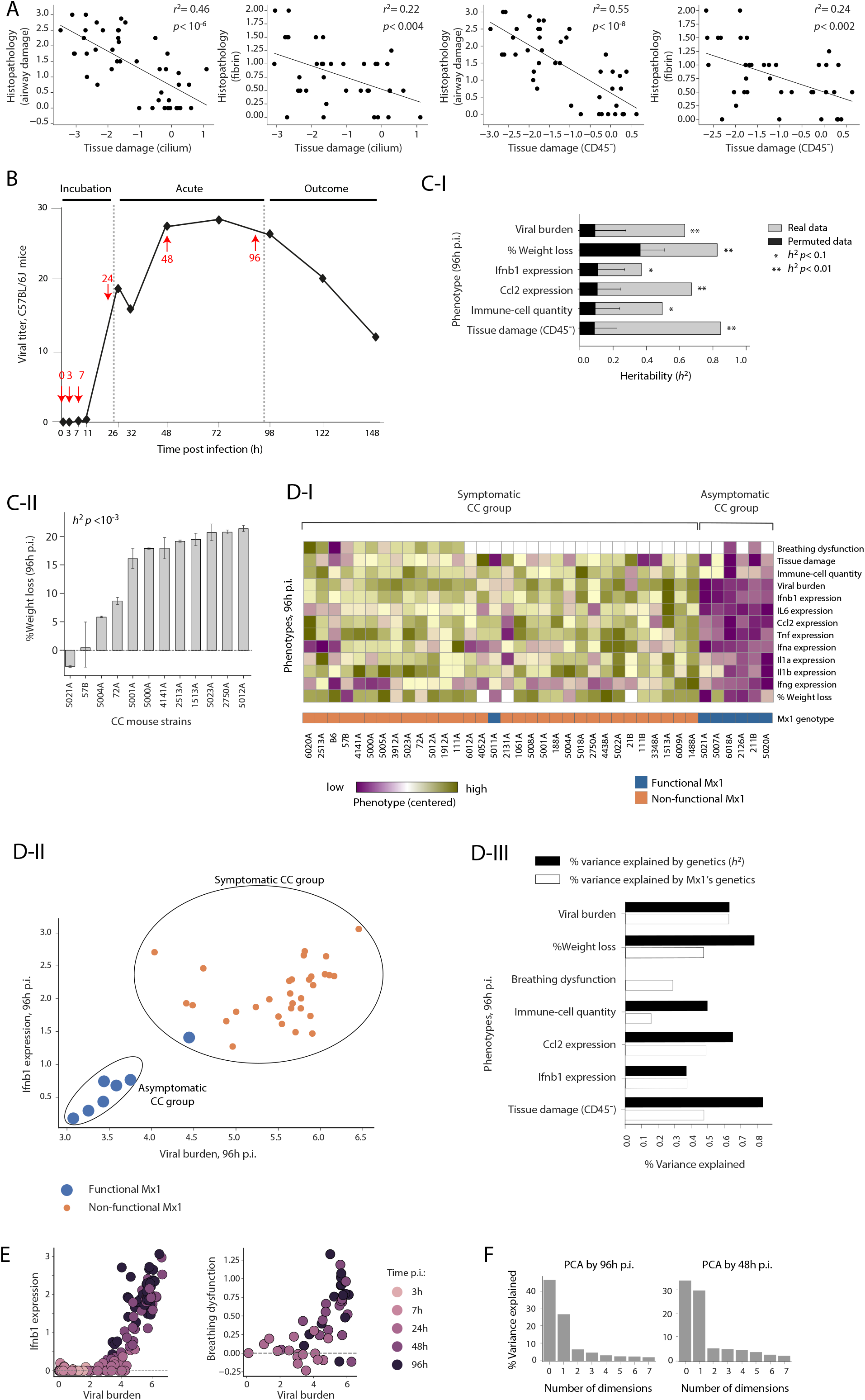
Diversity of IAV infection severity in various mouse strains. (**A**) Evaluation of the transcriptional signature of tissue damage. Shown are relations between the transcriptional tissue damage signature in lungs (*x* axis) and the pathophysiological status of lungs (y axis) across IAV-infected mice from 44 different genetic backgrounds at 4 days post infection (dots). The *x* axis presents the transcriptional signature of tissue damage, calculated based either on marker genes of CD45^−^ or ciliated cell types. This signature was used in the current study (e.g., Figure 1A). The *y* axis presents scoring of tissue damage in lungs based on histopathological examinations, including the protein level of fibrin and an assessment of the general airway damage. Phenotypic data from Ref. 20, and gene expression data was downloaded from GEO accession GSE30506. (**B**) Viral titers in lungs during IAV infection of the C57BL/6J mouse strain (adapted from Altboum et al., 2014). Red arrows indicate the time points of data collection in the current study. (**C**) Heritability of infection phenotypes. (I) Heritability (*h*^2^, defined as the percentage of between-strain variation from the total variation, *x* axis) of different disease phenotypes at 96h p.i. (*y* axis), either in real data (gray) or permuted data (black). Asterisks denote the significance of heritability. (II) The percentage of whole-body weight loss at 96h p.i. (*y* axis) across the CC mouse strains (*x* axis). Error bars were calculated using two mice per strain. (**D**) The baseline T levels are associated with late severity of infection in CC mice. (**D-I**) Characterization of symptomatic and asymptomatic CC strains, and the relations to genetic variation in the Mx1 gene. For each phenotype at 96h p.i. (rows) and each CC mouse strain (columns), shown is the measured phenotype (color coded, relative to the average across strains). The heatmap highlights a partition of individuals into two groups based on hierarchical clustering (symptomatic and asymptomatic). Variation in the Mx1 gene, whose function has a known, central influence on susceptibility to IAV infection (Ferris et al., 2013), is indicated (bottom). The matrix shows that Mx1-deficient mice tend to have a higher severity of infection (**D-II**) A scatter plot of viral burden (*x* axis) and *Ifnb1* expression (*y* axis) at 96h p.i., across strains (dots). Circles indicate the symptomatic and asymptomatic clusters from D-II. Color coding indicates strains that carry a functional (blue) or a non-functional (orange) Mx1 gene. The plot shows that Mx1-deficient mice tend to have a higher severity of infection. (**D-III**) Percentage of explained phenotypic diversity in the IAV-infected individuals. *Y* axis: phenotypes at 96h p.i. *X* axis: the percentage of phenotypic diversity that is explained by genetic background (i.e., the ‘inherited variation’, also referred to as ‘heritability’ (h^2^), black) and the percentage of phenotypic diversity that is explained by genetic variation in the Mx1 gene (white). Overall, the analysis emphasizes the presence of two phenotypic groups (symptomatic and asymptomatic strains), and further indicates that genetic variation in Mx1 explains much of the variation between these groups. (**E**) Relations between disease phenotypes and viral burden across the CC mouse cohort. The plots are shown as in Figure 1C but using all individuals. (**F**) Principal component analysis of the CC’s gene expression data at 96h p.i. (left) and 48h p.i. (right). For each principal component, shown is the total variation explained by the component.

**Supplementary Figure 2.**
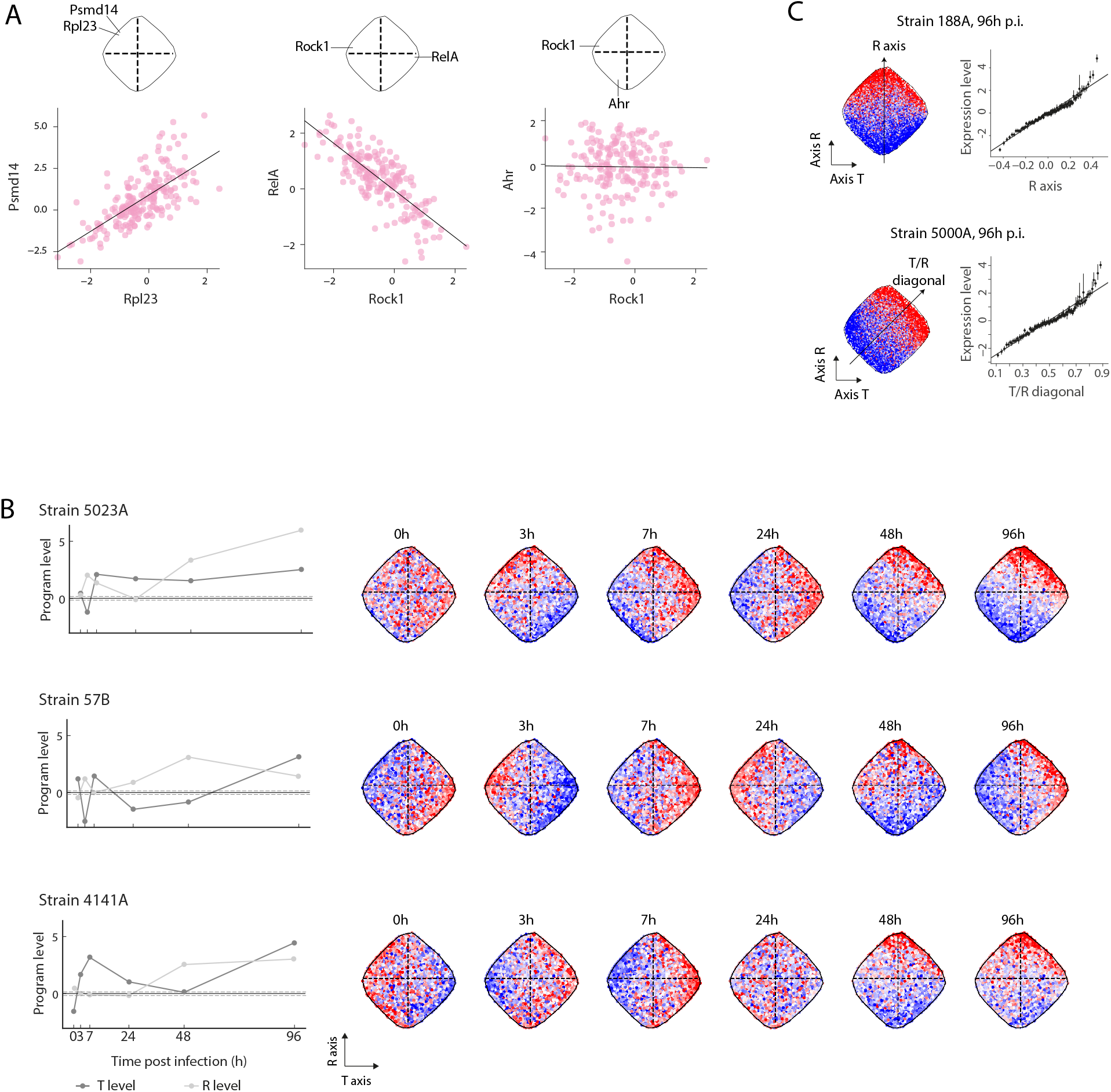
Basic characterization of the T/R model. (**A**) The similarity rule. Scatter plots for the relations between pairs of genes across all individual mice (all strains and time points). The positions of genes in the gene map (the map from Figure 1D-I) are indicated on top. The plots demonstrate the organization of the map: (i) nearby genes in the map are positively correlated (e.g., *Rpl23, Psmd14*), (ii) genes in opposite positions in the map are negatively correlated (e.g., *Rock1, RelA*), and (iii) other pairs of genes are either uncorrelated or weakly correlated (e.g., *Ahr, Rock1*). (**B**) Demonstration of T and R dynamics along the course of IAV infection, for three CC mouse strains (top, middle, and bottom). T and R levels are shown either by the calculated levels (left) or by coloring the gene map with the expression levels of each gene (right). Left: dashed lines indicate empirical *p* < 0.001 based on permutation analysis. (**C**) A nearly linear change in the expression of genes along the gradient. A detailed analysis of two individual mice at 96h p.i.: one from strain 188A (top panels) and the other from strain 5000A (bottom panels). For each individual, the left panel shows the gene map, color coded with its gene expression. The direction of the gene expression’s gradient is indicated as an arrow on top of the map. For each individual, the right panel presents a sliding window of expression levels along the direction of the indicated gradient.

**Supplementary Figure 3.**
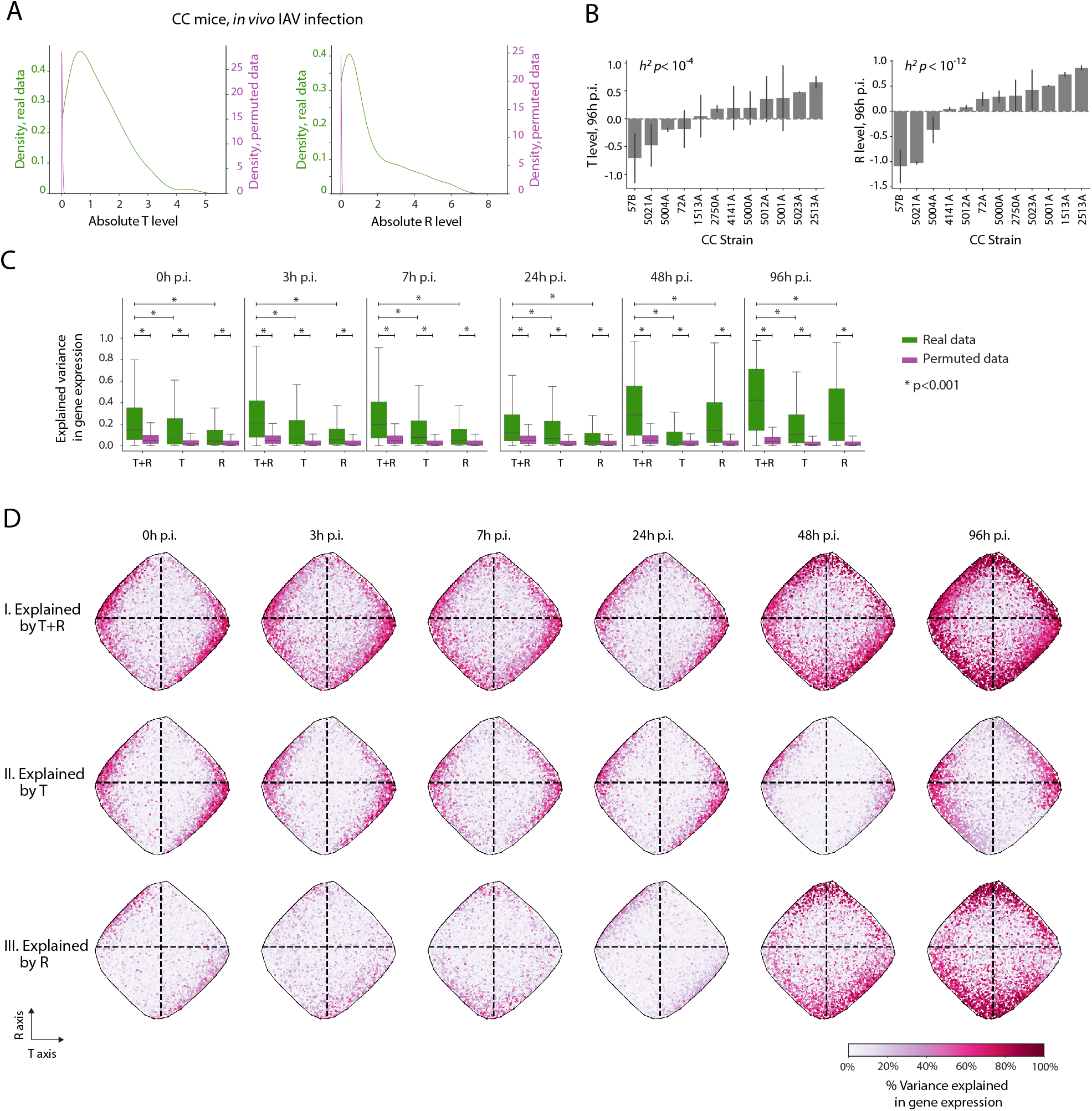
Corroboration of the R/ T model using the CC cohort. (A,B) Tests of explained variation. (**A**) Tests of significance. Distribution of absolute T and R levels (left and right, respectively) in the CC mice during *in vivo* IAV infection. T and R levels were calculated using the measured data (green) and permuted data (purple). The empirical p-value cutoff in Figure 1E and in **Figure S2B** is based on this analysis. (**B**) Tests of heritability. T and R levels (*y* axis) across 12 CC strains at 4 days p.i. (*x* axis). Error bars were calculated using two mice per strain. P-values of heritability are indicated on top. (**C**) The T and R scores explain substantial fractions of the variation in gene expression. Shown are distributions of the explained variation in gene expression across genes (*y* axis), calculated for different time points (plots, indicated on top), when using only T levels, only R levels, or both (*x* axis). For each model, explained variation was calculated using the measured data (green) and permuted data (purple). (**D**) Color coding of the gene map with the explained gene-expression variation. Each gene (a dot in the gene maps) is color-coded by its explained variation (white to pink scale). In each panel, the explained variation was calculated for a particular time point (left to right panels) using the combination of T and R (I), a T-only model (II), or an R-only model (III). The plots indicate that genes closer to the boundary of the map are better correlated with the T/R state. Overall, plots C and D indicate that the variation explained by an R+T model is larger than the variation explained by the R-only or T-only models.

**Supplementary Figure 4.**
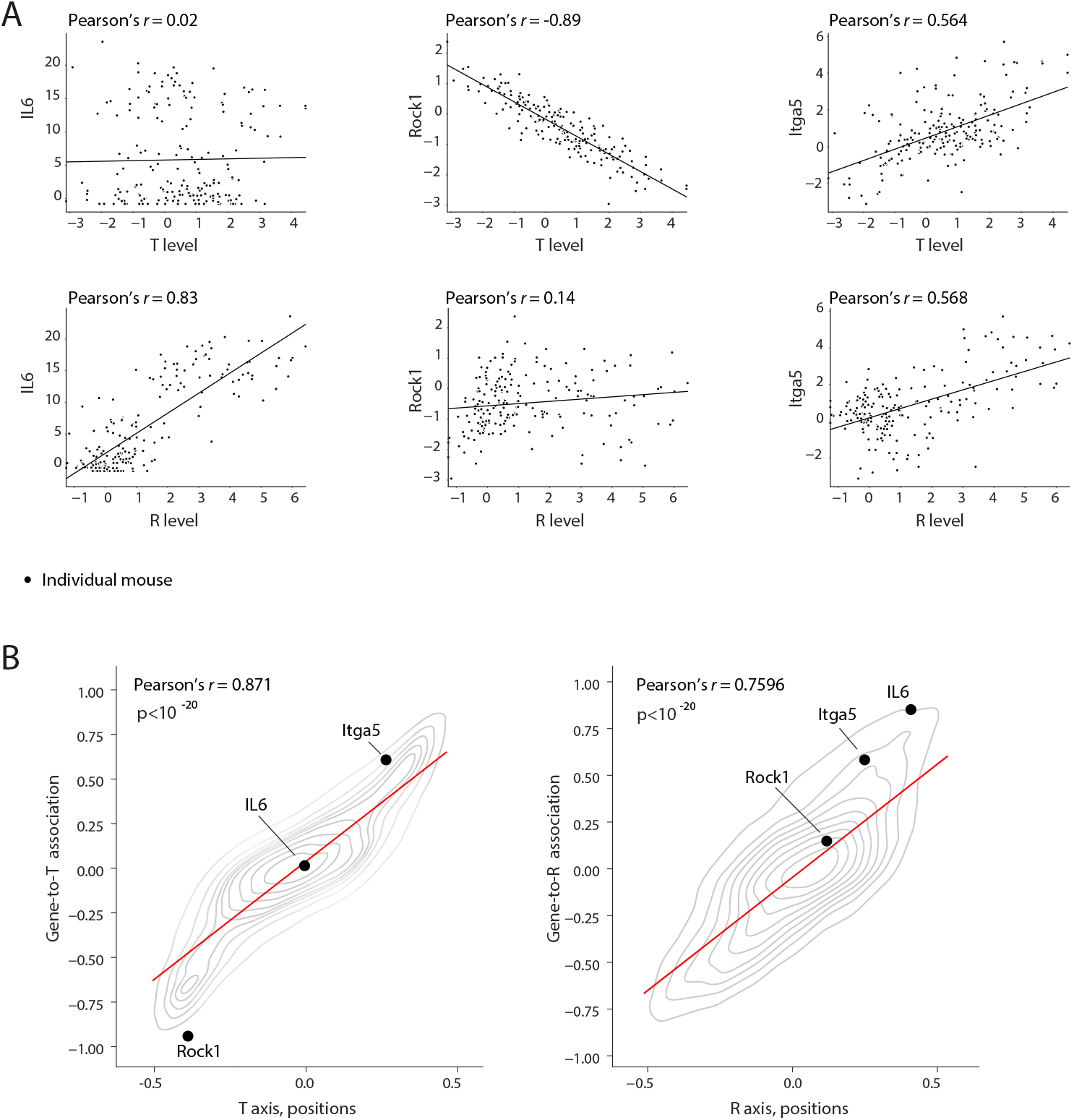
Gene-to-program associations and their consistency with the positions of genes in the map. (**A**) Examples of the relations between gene expression and the T/R programs. For each gene (a sub-panel), shown is a scatter plot of its expression level (*y* axis) and the levels of T (top) and R (bottom) (*x* axis) across all individuals (dots), including individuals of all time points and CC strains. The correlations, referred to as ‘associations’, are indicated. (**B**) A global view of the relations between the position of genes in the map and their associations with program levels. Relations between the position of genes in the map (*x* axis) and their gene-program associations (*y* axis), visualized using two-dimensional kernel density estimates (KDE) plots. The red lines indicate the fitted linear regression line. Specific genes from panel A are indicated. The plots show that genes closer to the end of an axis in the map are better correlated (or better anti-correlated) with the respective program.

**Supplementary Figure 5.**
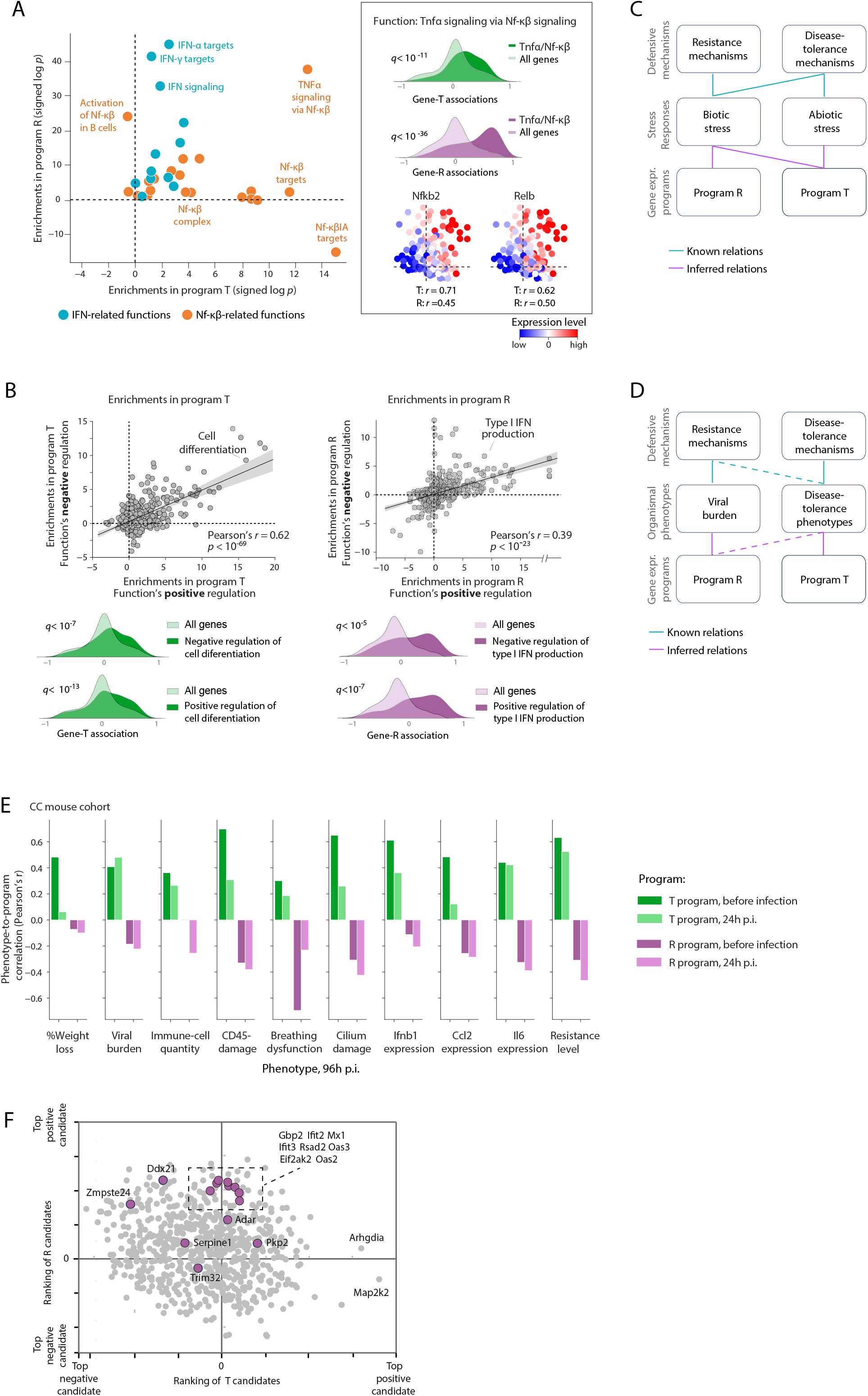
Functional characterization of the T and R programs. (**A**) Enrichments of NFκB-related functions (orange, e.g., ‘TNF-mediated NF-κB signaling’) and interferon-related functions (light blue, e.g., ‘type I IFN signaling’) with programs T and R, presented as in Figure 2A,B. (**B**) Coordination of positive and negative factors. Top: Enrichments of positive regulators (*x* axis) and negative regulators (*y* axis) of each functional category (dots) with program T (left panel) and program R (right panel). Bottom: Examples of specific enrichments, presented as in Figure 2B. (**C**) An overview of the relations between defensive mechanisms (top), response to stressors (middle), and the gene-expression T/R programs (bottom). Top versus middle layers: The relations between the top and middle layers (blue lines) reflect the well-established role of defensive mechanisms in biotic and abiotic responses. Bottom versus middle layers: The relations between the middle and bottom layers (purple lines) present the inferred associations in Figure 2C. (**D**) An overview of the relations between defensive mechanisms (top), organismal phenotypes during infection (middle), and the gene-expression T/R programs (bottom). Top versus middle layers: The relations between the top and middle layers (blue lines) reflect the well-established link between the defensive resistance strategy and the pathogen burden, as well as the role of both disease tolerance and resistance mechanisms in determination of disease tolerance phenotype. As in some systems resistance mechanisms do not have contribution to disease tolerance phenotypes, this relation is shown with a dashed line. Bottom versus middle layers: The relations between the middle and bottom layers (purple lines) presents the inferred associations in Figure 2D. As program R is associated with disease tolerance phenotypes only in H1N1 infection but not H3N2 infection (Figure 2D-II), this relation is shown with a dashed line. Overall, the illustration shows that programs T and R fit the signature of disease-tolerance and resistance mechanisms, respectively. (**E**) Pearson’s correlations between baseline/early program levels and late IAV infection severity phenotypes, across the Mx1-deficient CC strains (*y* axis). Disease phenotypes (at 96h p.i.) are indicated (*x* axis). Program levels are either before IAV infection or at 24h p.i. (color-coded). The plot shows that for all symptoms, the baseline T levels are negatively associated with late severity of infection. The baseline R levels have an opposite (and a much lower) effect. (**F**) Ranking of perturbation candidates by their averaged T/R-associations (shown as in Figure 5A, but with directions). All validated IAV restriction factors, as reviewed in Ref. 37, are highlighted in purple. This plot provides additional support for the hypothesis that program R is linked to resistance functions.

**Supplementary Figure 6.**
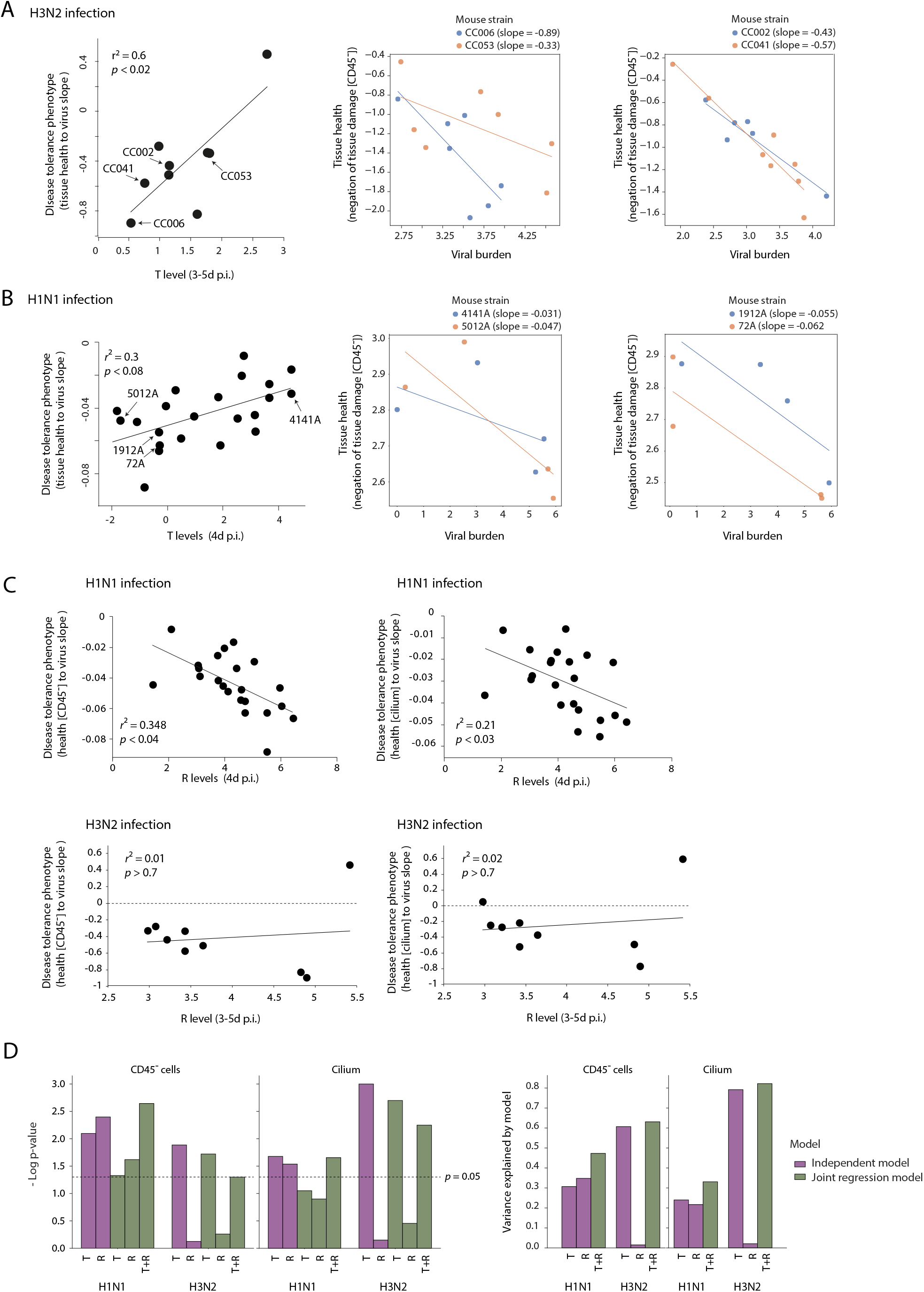
Relations of programs T and R with phenotypic diversity in disease tolerance. (**A, B**) Left: Relations of program T (x-axis) with the measure of disease tolerance phenotype (y-axis) across CC strains (dots). The plots are the same as in Figure 2D-II with additional indications of specific mouse strains. Right: Examples for the calculation of ‘disease tolerance phenotype’ using the reaction norms approach (Refs. 23, 25, 54). Shown are scatter plots for the relations between the health and viral burden in each mouse individual. For each mouse strain, the metric of ‘disease tolerance phenotype’ is calculated as the slope of the regression line across all individuals from this strain. The calculation of ‘disease tolerance phenotype’ (that is, the slopes) are shown for those particular strains that are indicated in the panel on the left side. A: H3N2 IAV infection (data from GSE136748); B: H1N1 IAV infection (dataset of this study). (**C**) Relations of program R (x-axis) with the measure of disease tolerance phenotype (y-axis) across CC strains (dots). Plots are shown as in Figure 2D-II but for the R program. (**D**) Summary of the relations between programs T/R and the phenotypic diversity in disease tolerance. Shown are the percentage of explained variation and the minus log p-value for the regression model in which the explained variable is the ‘disease tolerance phenotype’ and the explaining variable is the program level (several such regression models are demonstrated in panels A and C). The ‘independence model’ (purple) is a simple regression model in which the explaining variable is either program R or program T. The ‘joint model’ (green) is a multiple regression model in which both T and R are used as the explaining variables. Disease tolerance phenotypes were calculated using two alternative ‘health’ measures: (i) the negation of tissue damage, calculated using markers of CD45^−^ cells, and (ii) the negation of tissue damage, calculated using cilium markers (indicated on top). The plot demonstrates that program T is consistently associated with the phenotypic diversity in disease tolerance.

**Supplementary Figure 7.**
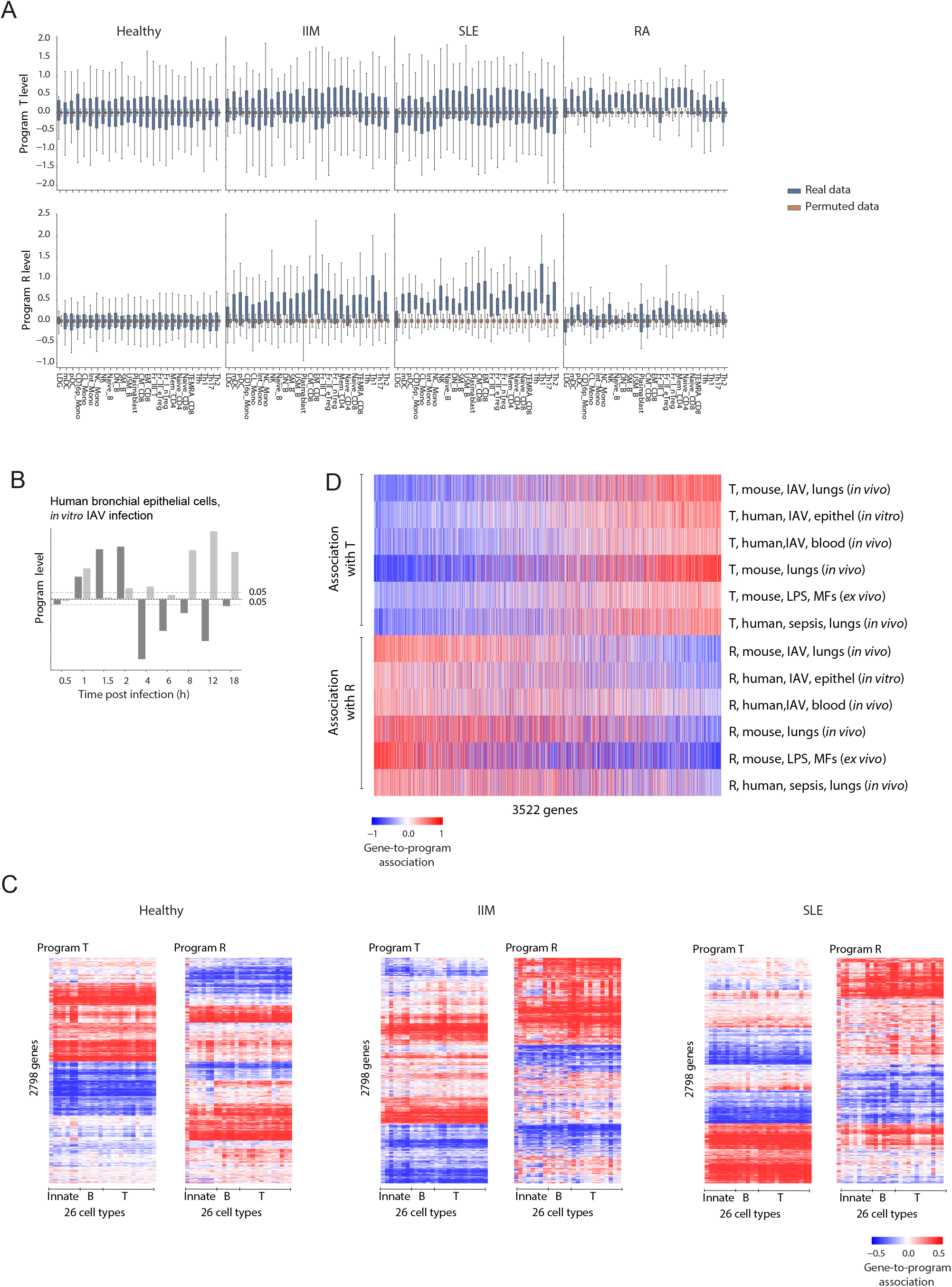
Programs T and R are relevant in all cell types under study. (**A**) Inter-individual variation in T (top) and R (bottom) levels (*y* axis), in each cell type and disease (*x* axis). Presented are the original and permutation-based levels. (**B**) Changes of T and R levels in primary human bronchial epithelial cells during the response to IAV infection (Ref. 27). The dynamics of T/R responses are in agreement with the dynamics of lung responses during the incubation and early acute phases of IAV infection. (**C**) Consistency of each program in all cell types under study. Shown are the gene-to-program associations (color-coded) for each gene (row) and each program (T/R: left/right), calculated based on transcription profiles in each cell type (column). Each matric shows associations that were calculated based on three different cohorts: associations were calculated based on data from healthy subjects (left), IIM patients (middle) and SLE patients (right). Figure 3A presents the averaging of associations across all five autoimmune-disease cohorts and the healthy cohort. (**D**) Consistency of gene-to-program associations across additional datasets. The heatmap presents associations (color-coded) between each gene (column) and the R or T level, using data from various independent datasets (rows, see **Methods** for source data). The heatmap highlights a consistency of associations between genes and the T/R levels across datasets.

**Supplementary Figure 8.**
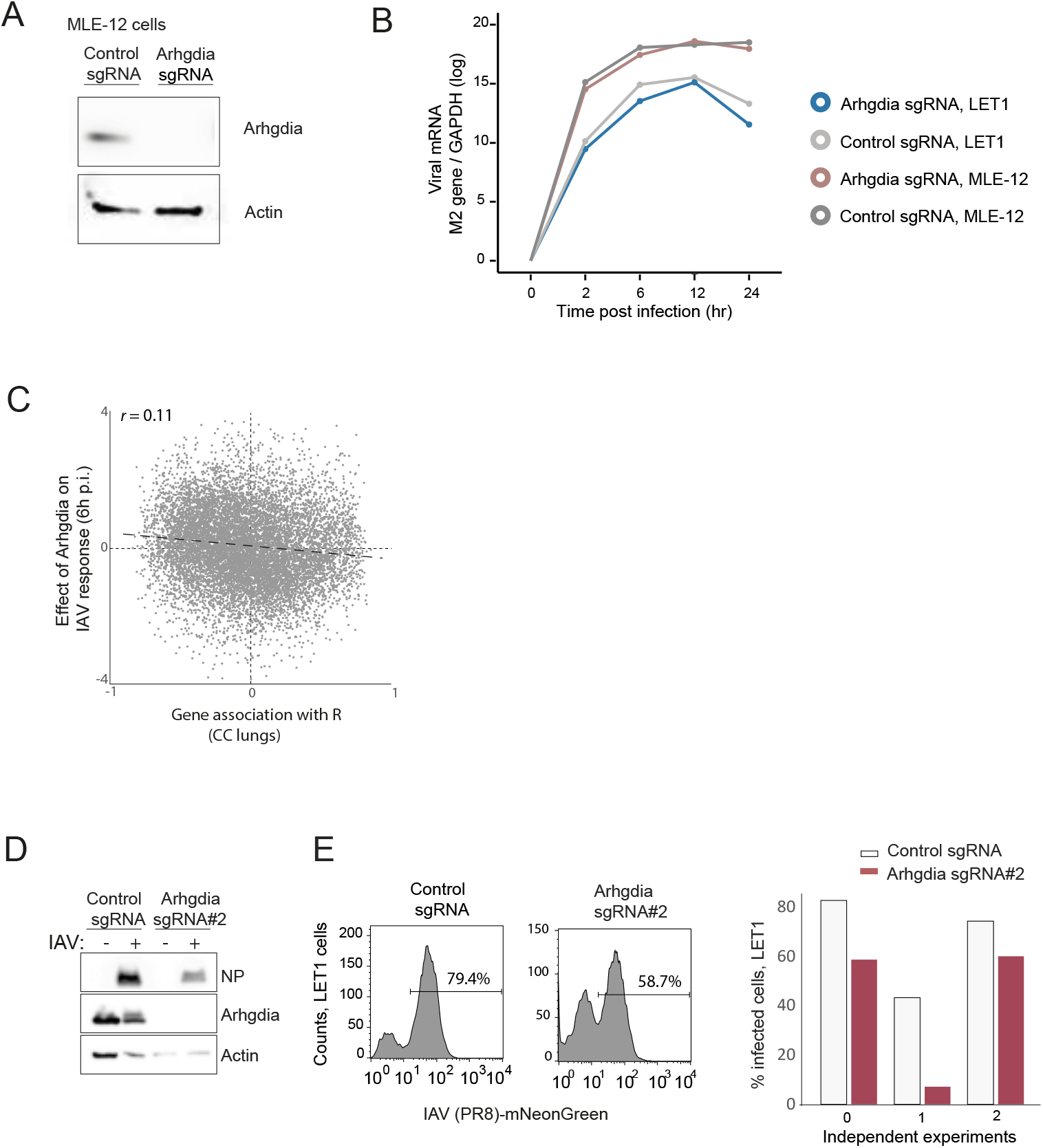
Arhgdia – additional data. (**A**) Knock out of Arhgdia using CRISPR-Cas9 editing in MLE-12 cells. Shown as in Figure 5B but for MLE-12 cells. (**B**) Viral RNA levels in cells expressing, or not, Arhgdia. The viral RNA levels were determined by qRT-PCR, using primers specific for viral M2 gene and cellular GAPDH, respectively. (**C**) Comparison between the signatures of program R and Arhgdia. For each gene (a dot), shown is its association with R levels (using lung data across CC mice; *x* axis) compared to the effect of Arhgdia on this gene (differential expression in LET1 cells: control 6h at p.i. versus Arhgdia-deletion at 6h p.i., *y* axis). We note that the similarity between the signatures of Arhgdia and program T (Figure 5C) was much stronger. (**D,E**) Reproducibility of the effect of Arhgdia on viral burden. Data shown as in Figure 5B,E but for a different sgRNA clone (denoted dgRNA#2).

**Supplementary Table 1**. **Measured phenotypes in the CC cohort**. For each CC strain (column 1), reported are the standard UNC identifier (column 2), genetics of the Mx1 gene (column 3), and the group of susceptibility to IAV infection (column 4, based on the analysis of disease severity in **Figure S1D**). Columns 5-16 report the T and R levels of each individual (i.e., for each strain and time point). Columns 17-55 report additional disease phenotypes in each individual mouse. Columns 5-55: time points following IAV infection (baseline: before IAV infection). Top 33 strains: most time points were measured. Bottom 6 strains: only late time points were measured.

**Supplementary Table 2. The map.** For each gene (column 1), reported are its coordinates in the map (columns 2-3) and its associations with R levels and T levels (columns 4-5) across the CC IAV cohort. The coordinates in the map allow the calculation of the quantitative levels of T and R in each given sample (using deconvolution as formalized in Eq. 1).

**Supplementary Table 3. Correlations of disease phenotypes with baseline T and R levels across the BXD mouse strains**. A collection of autoimmunity, infectious disease and liver injury phenotypes across the BXD mouse lines. For each category of phenotypes (column 1), included are details about the phenotype, correlations of the phenotype with T and R levels in resting or activated MFs. Specific transformations and references for categorizations are also reported. Included are all phenotypes in the GeneNetwork database (Mulligan et al. 2017) for which (*i*) the phenotype falls into one of the categories in column 1, and (*ii*) the phenotypes was measured in at least 5 BXD strains for which transcriptomes in resting/activated MFs were measured (MF transcriptomes from GSE38705).

**Supplementary Table 4. Ranking of T and R candidate regulators.** Reported is the ranking score of regulators. Positive/negative for regulators that are positively/negatively associated with T or R levels.

**Supplementary Table 5. Percentage of inherited variation in disease severity that is explained by the baseline T and R levels**. For each phenotype of the CC mice at 96h post IAV infection (column 1), reported are: (i) the percentage of total variation that is explained by variation in baseline (before infection) T/R levels (columns 2,4), and (ii) the percentage of inherited variation that is explained by variation in baseline (before infection) T/R levels (columns 3,5). Calculations are based on inherited variation (heritability) values that are reported in **Figure S1C**.

## Acknowledgments

This work was supported by the European Research Council (ERC 637885) and by the European Union Horizon 2020 under grant agreement No. 847422. GY, OC, NPY, YS, AF and RB were supported by ERC 637885. GY, YS, AF and RB were supported by the Edmond J Safra Center for Bioinformatics at Tel Aviv University. IG-V is a Faculty Fellow of the Edmond J Safra Center for Bioinformatics at Tel Aviv University. IGV, EB, OC, GY and NPY are listed as inventors on a filed provisional patent for blood markers of disease tolerance and resistance states. All other authors do not disclose any conflicts of interest.

